# Decoupling of inter-regional functional connectivity and regional neural activity in Alzheimer Disease

**DOI:** 10.1101/642629

**Authors:** Somayeh Maleki-Balajoo, Farzaneh Rahmani, Reza Khosrowabadi, Chun Meng, Timo Grimmer, Alexander Drzezga, Mojtaba Zarei, Christian Sorg, Masoud Tahmasian

## Abstract

Alzheimer’s disease (AD) and mild cognitive impairment (MCI) are characterized by aberrant regional neural activity and disrupted inter-regional functional connectivity (FC). It is, however, poorly understood how changes in regional neural activity and inter-regional FC interact in AD and MCI. Here, we investigated the link between regional neural activity and nodal topological measures of FC through simultaneous PET/MR measurement in 20 patients with MCI, 33 patients with AD, and 26 healthy individuals. First, we assessed regional glucose metabolism identified through FDG-PET (rFDG) (as a proxy of regional neural activity), and regional FC topology through clustering coefficient (CC) and degree centrality (DC) (as surrogates of local segregation and global connectivity, respectively). Next, we examined the potential moderating effect of disease status (AD or MCI) on the link between rFDG and FC topology using hierarchical moderated multiple regression analysis. Alterations in rFDG, CC, and DC were widespread in patients, and AD alters physiological coupling between regional metabolism and functional connectivity particularly in the inferior temporal gyus and supplementary motor areas. While rFDG correlated with CC in healthy subjects, this correlation was lost in AD patients. We suggest that AD pathology decouples the normal association between regional neural activity and functional segregation.

## INTRODUCTION

Alzheimer disease (AD) is characterized by distinct patterns of neuronal loss leading to reduced overall neural activity, which can be indirectly quantified through fluorodeoxyglucose-positron emission tomography (FDG-PET) (Minoshima, et al., 1995; Mosconi, 2013). It has been well-documented that regional glucose metabolism (rFDG) is correlated – to distinct degrees - with cortical atrophy, beta-amyloid and tau protein accumulation, which are hallmarks of AD pathology (Bischof, et al., 2016; Chetelat, et al., 2008; Drzezga, et al., 2008). Functional connectivity (FC) is quantified as the magnitude of functional co-activation between distinct brain regions’ neural activity, typically measured by correlated blood oxygenation fluctuations of distinct regions of resting-state functional MRI (rs-fMRI). As 80% of neuronal metabolic activity is dedicated to synaptic signalling (Attwell and Laughlin, 2001; Dienel, 2019), FC might also demonstrate a conjugate increase or decrease in neural activity between brain regions. Indeed, studies have pointed out that a common neural substrate underlies rFDG and FC in healthy individuals and in AD patients (Marchitelli, et al., 2018; Passow, et al., 2015; Riedl, et al., 2014; Savio, et al., 2017).

Alterations in resting-state FC patterns has been observed along the trajectory of AD (Badhwar, et al., 2017; Greicius, et al., 2004; Sorg, et al., 2007). Topological features of these FC patterns can be modeled through graph theory analyses, where brain regions are considered as “nodes”, and the connectivity between them as “edges” of the graph (Latora and Marchiori, 2001; Rubinov and Sporns, 2010). Clustering coefficient (CC) and degree centrality (DC) are two commonly used nodal topological metrics, representing local segregation and centrality of individual nodes, respectively (Masuda, et al., 2018). In particular, CC quantifies how well a certain node has formed locally-segregated processing modules around itself (i.e., how efficient a node is clustering with other nodes). Meanwhile, DC gives a measure of the importance/centrality of a node in terms of interacting with other nodes within and outside the regional clusters (i.e., how well it can preserve regional segregation and global integration of the network at the same time) (Rubinov and Sporns, 2010).

A robust link between FC and metabolic brain networks has been identified in healthy brain (Riedl, et al., 2014). This association, however, has been shown to be disturbed in AD, where rFDG no longer correlates with regional functional activity in target AD regions in posterior associational cortices (Marchitelli, et al., 2018). It has been suggested that the local amyloid beta pathology might be responsible for decoupling between rFDG and inter-regional FC in the default mode network (DMN) in AD patients (Scherr, et al., 2018). Whether and how AD pathology affects the link between rFDG and FC topological characteristics is poorly understood. Here, we applied a descriptive-analytic approach to address these questions based on simultaneous acquisition of FDG-PET and rs-fMRI, using a hybrid PET/MR scanner in patients with AD and mild cognitive impairment (MCI), as well as in healthy subjects.

## METHODS

### Participants

Thirty three patients with mild AD-dementia, 20 patients with MCI and 26 healthy subjects were recruited in this cross-sectional study. Patients were randomly selected from outpatient memory clinic of the Department of Psychiatry and Psychotherapy of Klinikum rechts der Isar of Technical University of Munich, School of Medicine (TUM). The AD or MCI status of patients was determined using Clinical Dementia Rating (CDR) (Morris, 1993) and neuropsychological testing batteries based on criteria established by Consortium to Establish a Registry for Alzheimer’s disease CERAD (Welsh, et al., 1994). MCI patients had a CDR global of 0.5 and evidence of cognitive impairments in neuropsychological testing with intact activities of daily living (Gauthier, et al., 2006), while patients with AD fulfilled criteria for mild AD based on National Institute on Aging-Alzheimer’s Association criteria (McKhann, et al., 2011). A summary of subjects’ demographic information is provided in Table 1. Twenty-six healthy subjects were also recruited through word-of-mouth advertising in Munich. The study was approved by the university ethics committee (TUM) in line with the institute’s Human Research Committee guidelines and conformed to standards of the declaration of Helsinki. All participants provided informed consent after being given proper explanation about the goal and protocol of the study.

**Table 1.** Demographic and clinical data of participants.

|  | **HC (n=26)** | **MCI (n=19)** | **AD (n=33)** | **p-Value** |
| --- | --- | --- | --- | --- |
| **Age, year** (*mean ± SD*) | 55.58±10.21 | 68.05±9.82 | 71.06±8.78 | <0.001 |
| **Sex** (*female*) | 9 | 12 | 14 | 0.157 |
| **MMSE score** (*mean ± SD*) | 29.51±1.02 | 26.94±2.16 | 22.71±4.18 | <0.001 |
| **CREAD NAB total**<br>(*mean ± SD*) | 86.10±8.13 | 68.75±10.41 | 55.13±11.95 | <0.001 |

Abbreviations: AD: Alzheimer disease; HC: healthy controls; MCI: mild cognitive impairment; MMSE: Mini-Mental State Examination; CERAD: Consortium to Establish a Registry for Alzheimer's Disease neuropsychological assessment battery. Analysis of variance, $p < 0.05$ as threshold of significance, except of sex (Kruskal-Wallis test).

### Data acquisition and preprocessing

Imaging data acquisition included structural MRI rs-fMRI and FDG-PET which were simultaneously acquired on a Biograph mMR scanner (Siemens, Erlangen, Germany). MR images were preprocessed using Statistical Parametric Mapping (SPM12) and the Data Processing Assistant for Resting-State fMRI toolbox (DPARSF) (Yan, et al., 2016) as previously performed (Riedl, et al., 2014; Scherr, et al., 2019; Tahmasian, et al., 2015; Tahmasian, et al., 2016). In order to perform multimodal preprocessing, we first co-registered mean FDG-PET image to the T1-weighted image using rigid-body transformation. Both images were then co-registered to the mean rs-fMRI image. We performed the following steps for preprocessing of the rs-fMRI images: removing the first 3 volumes, realignment, normalization and smoothing (Satterthwaite, et al., 2013; Tahmasian, et al., 2015; Tahmasian, et al., 2016), detrending, band pass filtering (0.01-0.1), and regressing out the nuisance variables including motion parameters with the Friston-24 model, the white matter and mean cerebrospinal fluid signal as previously suggested (Satterthwaite, et al., 2013). Partial volume correction of FDG-PET image was carried out using PMOD software (PMOD Technologies Ltd., Adliswil, Switzerland), normalization and smoothing as described previously (Tahmasian, et al., 2015; Tahmasian, et al., 2016). Additional details regarding the data acquisition, movement corrections, and preprocessing steps are provided in the Supporting Information.

### FDG-PET data analysis

The mean glucose uptake values of all voxels within the 112 regions of interest (ROIs), defined based on Harvard-Oxford atlas (Keuken, et al., 2014), were extracted from each subject-specific’s preprocessed PET images. We then normalized the mean glucose uptake value of each ROI by dividing it to the whole-brain glucose uptake value (Cranston, et al., 2001; Reed, et al., 1999). Analysis of variance (ANOVA) was then applied to identify group differences in rFDG of those 112 ROIs. The results were corrected for multiple comparisons using N-region statistical comparison as described previously (Lynall, et al., 2010; Meng, et al., 2014). The significance threshold was calculated using 1/number of ROIs, as: 1/112=0.009. Finally, post-hoc analyses (permutation test, 100,000 iterations, p < 0.05) was applied to identify significant rFDG differences between each pair of groups.

### rs-fMRI data analysis

#### Graph construction

Mean blood-oxygen-level dependent (BOLD) signal of each ROI was calculated. FC was defined as the functional links between the mean BOLD signals of every possible pair of ROIs (112 cortical and subcortical regions for each subject). Regional FC patterns were then constructed for 112 ROIs and presented based on the graph theory (Bullmore and Sporns, 2009), where brain regions served as nodes, and significant FC between them using absolute values of Pearson’s correlation coefficients, as edges of the graph (Friston, et al., 1993). Finally, a 112×112 symmetric undirected weighted matrix representing individual whole brain inter-regional FC was built for each subject.

From a topological perspective, each individual has a different topology in terms of connection density of brain regions. The connection density in a graph is defined as the ratio of the number of existing edges to the all possible edges (Bullmore and Sporns, 2012). Difference in connection density, in turn, influences most of the extracted topological metrics within a graph (Kaiser, 2011). Thus, it is necessary to implement a matching strategy between FC graphs prior to statistical analyses between the three groups (van Wijk, 2010). Accordingly, we generated the whole brain weighted FC metrics for the density range from 0.01–0.40 (with intervals of 0.01), as suggested previously (Sadeghi, et al., 2017). We then characterized the inter-regional FC’s organization of brain regions in terms of nodal CC (as a proxy of local FC) (van den Heuvel and Hulshoff Pol, 2010) and DC (as a proxy of global FC) (Sporns, 2013). To assess nodal topological metrics, we selected a wide density range and the integral values of metrics at the selected range were used in group comparison (Gong, et al., 2009). The mathematical definitions of these nodal topology metrics are available in the Supporting Information.

#### Nodal topological metrics analysis

The Brain Connectivity Toolbox (Rubinov and Sporns, 2010) was used for nodal analysis of weighted graph. The ANOVA model was implemented to perform group comparisons of nodal topological measures, i.e. CC and DC. The results were corrected for false positive results using the N-region statistical comparison as described above (Lynall, et al., 2010; Meng, et al., 2014). Post-hoc analyses were then conducted to identify group differences in CC and DC (permutation test, 100,000 iterations, p < 0.05).

### Cross-modality analysis

In order to explore the relationship between rFDG and FC topology metrics and investigate the role of MCI or AD in the relationship between rFDG and regional FC topology, we performed hierarchical moderated multiple regression (HMMR) analysis (see Supporting Information for more details) in the ROIs that AD pathophysiology affected both rFDG and inter-regional FC topology. Those ROIs were simply defined by overlapping the results of group comparisons of rFDG and CC or DC (AD vs. HC, AD vs. MCI, and MCI vs. HC). The potential moderating effect of MCI or AD on the link between rFDG and inter-regional FC topology, was examined using HMMR analysis in those overlapped brain regions.

We performed the HMMR analysis in two steps: First, rFDG and groups index (i.e. AD, MCI or HC) were entered as main predictors, while other covariates (i.e., age and sex) were added as covariates of no-interest, to a model to predict topological metrics (i.e. CC and DC). Second, the interaction as the product term of the main predictors, “rFDG × groups index” was added to the prediction model. To evaluate whether the interaction between rFDG and groups index was meaningful, we tested whether adding the “rFDG × groups index” increased the variance explained by the model in successive regression steps (Δ*R*^2^). When a statistically significant interaction emerged, it was interpreted according to the available guidelines (Aiken, 1991; Whisman and McClelland, 2005). Statistical Package for the Social Sciences V18 (SPSS Inc., Chicago, IL, USA) and Jamovi project (project, 2018) were used for HMMR analysis.

## RESULTS

### Glucose metabolism alterations along the trajectory of disease

Comparing rFDG between the three study groups revealed significantly reduced metabolism in various regions within the temporal lobe, angular gyri, precuneus, and lateral occipital regions in patients with MCI and AD compared to HC (Figure 1). Hypometabolism was most prominent in AD with MCI patients adopting an intermediate position between AD and HC. We also demonstrated hypermetabolism in the right hippocampus, bilateral parahippocampal, precentral, fusiform and lingual gyri, as well as bilateral brain stem, right amygdala and right supplementary motor area (SMA) in AD patients compared to MCI and HC groups. More information on between group comparisons of rFDG are provided in the Supporting Information.

**Figure 1.**
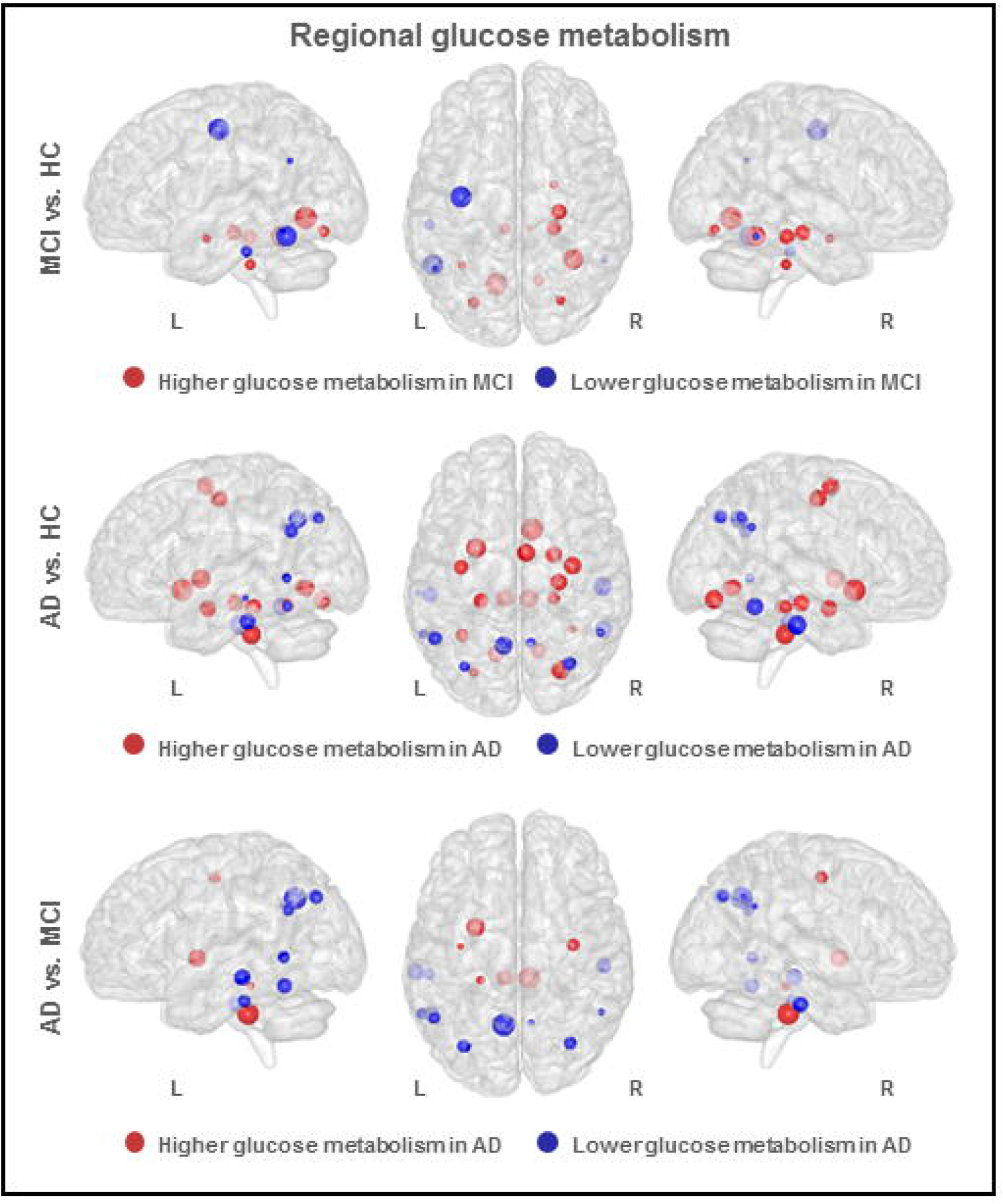
Analysis of variance on regional glucose metabolism for each region in Harvard-Oxford Atlas. Post-hoc test: permutation test (p < 0.05, 100,000 permutations). The size of each node corresponds to absolute mean difference values of regional glucose metabolism between each pairs of groups. AD: Alzheimer disease; HC: healthy controls; MCI: mild cognitive impairment; L: left; R: right.

### Disrupted local network topology along the trajectory of disease

Similar to rFDG, we found widespread changes in nodal topological metrics of inter-regional FC between each pair of study groups (Figure 2). CC was more extensively affected than DC in both patients groups, with progressive decrease starting from MCI in widespread cortical regions (Figure 2 and Table 2 in Supporting Information). Meanwhile, DC was increased in bilateral temporal fusiform gyri, right brain stem, and left pallidum (Figure 2B and Supporting Information).

**Table 2.** Overlapping brain regions determined in glucose metabolism analysis and inter-regional functional connectivity topology metrics (CC: clustering coefficient, DC: degree centrality)

|  | Brain regions | Regional glucose metabolism | Inter-regional functional connectivity topology metrics |  |
| --- | --- | --- | --- | --- |
|  |  |  | CC | DC |
| **MCI relative to HC** | *Precentral Gyrus (L) | ↓ | ↓ | - |
|  | *Inferior Temporal Gyrus, temporooccipital (L) | ↓ | ↓ | - |
|  | Temporal Occipital Fusiform Cortex (L) | ↑ | ↓ | - |
|  | Temporal Occipital Fusiform Cortex (R) | ↑ | ↓ | - |
|  | Occipital Fusiform Gyrus (L) | ↑ | ↓ | - |
|  | Occipital Fusiform Gyrus (R) | ↑ | ↓ | - |
| **AD relative to HC** | Precentral Gyrus (L) | ↑ | ↓ | ↓ |
|  | Precentral Gyrus (R) | ↑ | ↓ | ↓ |
|  | *Middle Temporal Gyrus, posterior (L) | ↓ | ↓ | - |
|  | *Middle Temporal Gyrus, temporooccipital (L) | ↓ | ↓ | - |
|  | *Inferior Temporal Gyrus, temporooccipital (L) | ↓ | ↓ | - |
|  | *Inferior Temporal Gyrus, temporooccipital (R) | ↓ | ↓ | - |
|  | *Angular Gyrus (R) | ↓ | ↓ | - |
|  | *Lateral Occipital Cortex, superior (R) | ↓ | ↓ | ↓ |
|  | Supplementary Motor Area (R) | ↑ | ↓ | - |
|  | *Precuneous Cortex (R) | ↓ | ↓ | - |
|  | Temporal Occipital Fusiform Cortex (L) | ↑ | ↓ | - |
|  | Temporal Occipital Fusiform Cortex (R) | ↑ | ↓ | - |
|  | Occipital Fusiform Gyrus (L) | ↑ | ↓ | - |
|  | Occipital Fusiform Gyrus (R) | ↑ | ↓ | - |
|  | *Brain-Stem(R) | ↑ | - | ↑ |
| **AD relative to MCI** | Precentral Gyrus (L) | ↑ | - | ↓ |
|  | Precentral Gyrus (R) | ↑ | ↓ | ↓ |
|  | *Middle Temporal Gyrus, posterior (L) | ↓ | ↓ | - |
|  | *Angular Gyrus (R) | ↓ | ↓ | - |
|  | *Lateral Occipital Cortex, superior (R) | ↓ | ↓ | - |
|  | *Brain-Stem(R) | ↑ | - | ↑ |

\* The direction of group differences in regional glucose metabolism and inter-regional functional connectivity topology metrics is congruent.
AD: Alzheimer disease; HC: healthy controls; MCI: mild cognitive impairment; L: left; R: right; ↑: increase; ↓: decrease.

**Figure 2.**
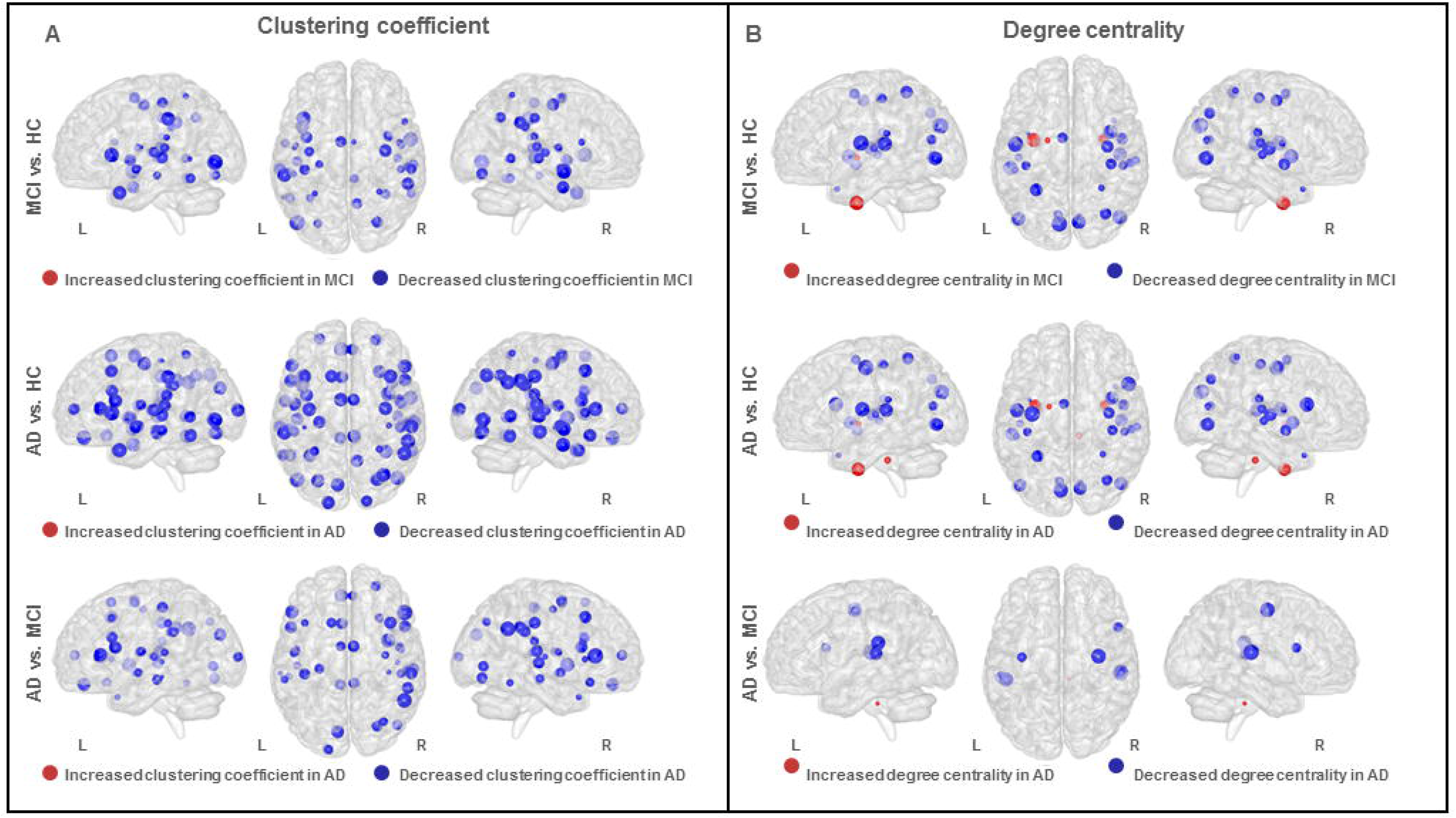
Analysis of variance on whole brain FC topology metrics for each region in Harvard-Oxford Atlas in A) clustering coefficient; B) degree centrality. Post-hoc test: permutation test (p < 0.05, 100,000 permutations). The size of each node corresponds to absolute mean difference values of clustering coefficient/degree centrality between each pairs of groups. FC: functional connectivity; AD: Alzheimer disease; HC: healthy controls; MCI: mild cognitive impairment; L: left; R: right.

### The effects of AD on the link between the regional glucose metabolism and the nodal topological measures

In order to investigate the potential effects of AD on the association between rFDG and topological metrics of inter-regional FC, we identified regions where rFDG changes and CC/DC changes overlapped (Figure 3 and Table 2). Hypometabolism was observed along with a reduction in CC in the bilateral middle temporal gyri, bilateral inferior temporal gyri-temporooccipital part (ITG), right angular, right lateral occipital and right precuneal cortices, while hypermetabolism corresponded to reduced CC or DC topological metrics in bilateral temporal occipital fusiform and occipital fusiform cortices, precentral gyri, and right SMA in patients groups compared to controls. The right brain stem showed an increase in both regional metabolism and DC in AD patients compared to MCI and HC.

**Figure 3.**
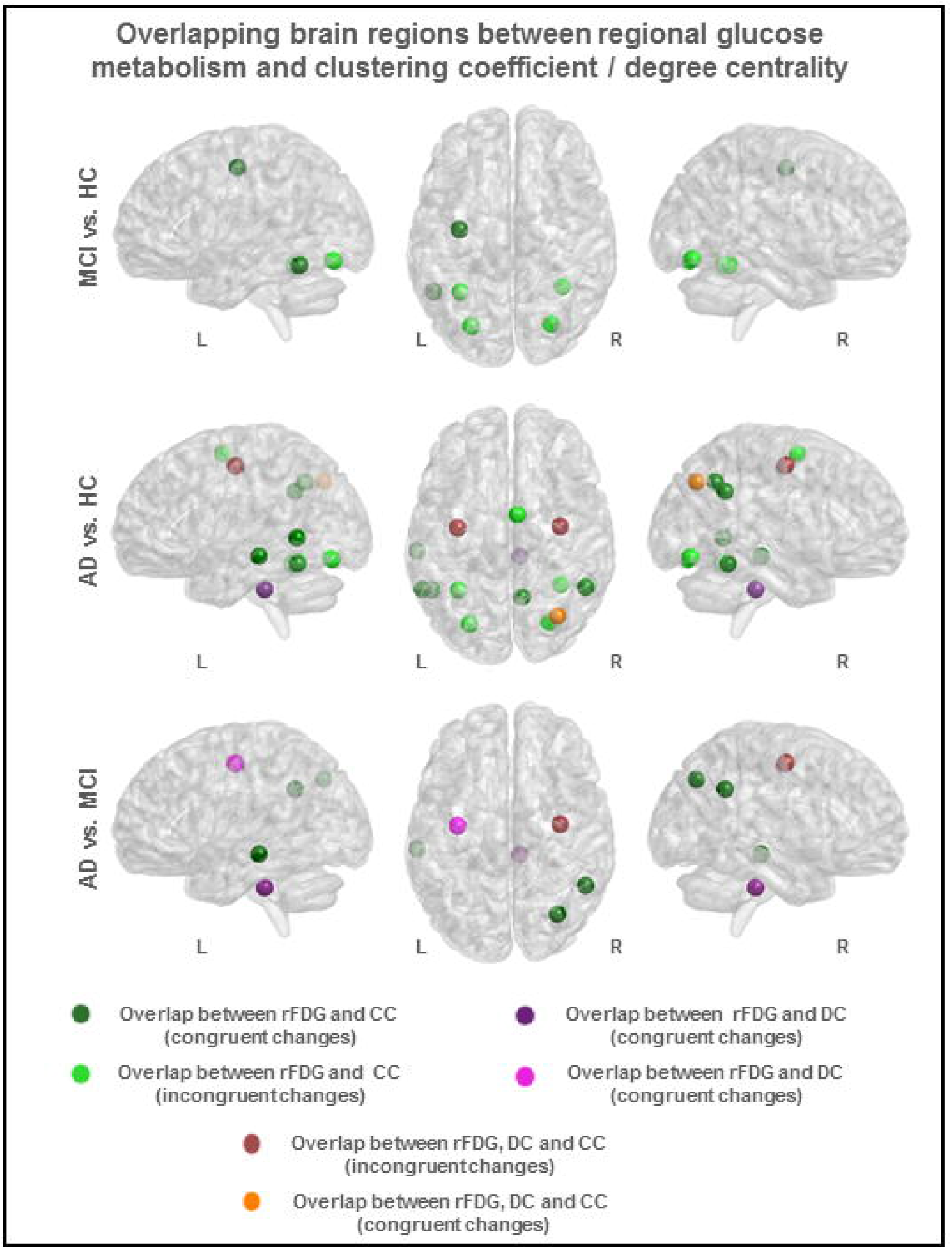
Overlapping brain regions between regional glucose metabolism and inter-regional FC topology metrics (clustering coefficient, and degree centrality). Congruent changes refer to similar direction of regional glucose metabolism and clustering coefficient/degree centrality between each pairs of groups. rFDG: regional glucose metabolism identified through FDG-PET; CC: clustering coefficient; DC: degree centrality; FC: functional connectivity; AD: Alzheimer disease; HC: healthy controls; MCI: mild cognitive impairment; L: left; R: right.

Adopting the HMMR analysis, we assessed whether rFDG and inter-regional FC topology interact in patients with AD and MCI and in HC. We demonstrated that a two-way interaction between “rFDG × groups index” in the following brain regions was able to significantly predict changes in CC in the right ITG (MNI coordinate: [19,39,29], *R*^2^ = 0.31 F (5,53) = 4.68, p_(model)_ = 0.001) and right SMA (MNI coordinate: [43,63,66], *R*^2^ = 0.5, F (5,53) = 10.71, p_(mode)l_ < 0.001) (Figure 4 A1 and B1). Adding the (rFDG × groups index) interaction, significantly improved the model in right ITG (Δ*R*^2^ = 0.08, p_(rFDG*group index)_ = 0.015) and in right SMA (Δ*R*^2^ = 0.046, p_(rFDG*group index)_ = 0.03). No significant model was able to predict changes in CC, when comparing AD vs. MCI or MCI vs. HC. In addition, the interaction between rFDG and groups index could not predict DC changes in any of the between group comparisons.

**Figure 4.**
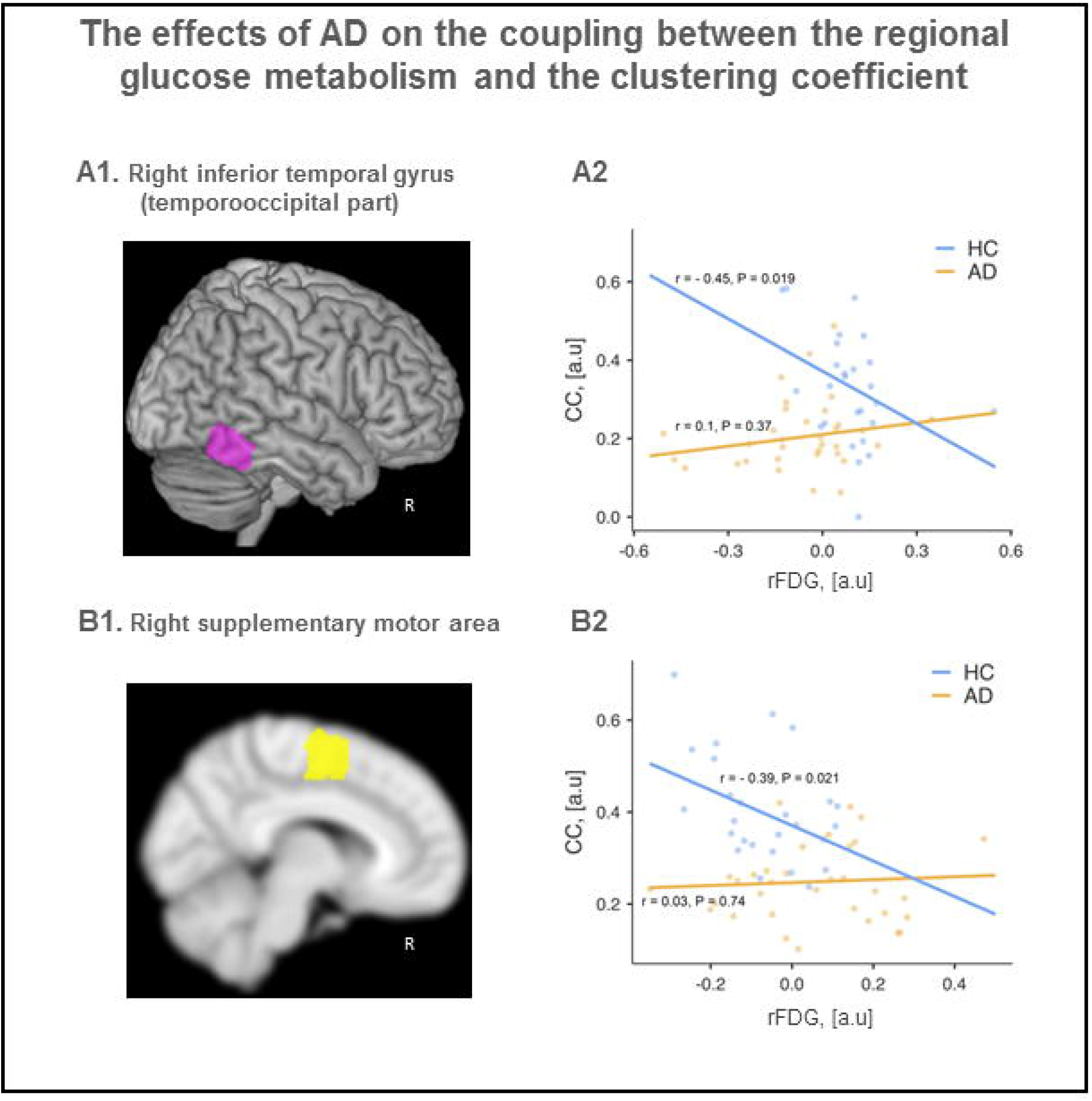
AD moderated the effect of regional glucose metabolism on predicting clustering coefficient. Results demonstrated that only in right ITG-temporooccipital part (MNI coordinate: [19,39,29], Figure 4 A1 and right SMA (MNI coordinate: [43,63,66], Figure 4 B1) defined by group) comparison between AD and HC, HMMR analysis predicted CC significantly with the two-way interaction (rFDG * groups index). Figure 4 A2 and B2 illustrated the effect of groups index defined as AD or HC on association between rFDG and CC in both right ITG and right SMA. In the right ITG, the effect of groups index as a moderator on relationship between rFDG and CC computed when moderator levels equals to HC is negative and strong, r = − 0.45, t (53) = − 2.42, p = 0.019. While AD was considered as the moderator levels, the effect becomes positive but weak and not significant, r = 0.1, t (53) = 0.91, p = 0.369 (Figure 4 A2). About right SMA, we also computed the relationship between rFDG and CC for different levels of the moderator (HC/AD) as well. Results showed when moderator levels equal to HC, the effect is negative, r = − 0.39, t (53) = − 2.37, p = 0.021. Finally, the effect calculated when we considered AD as the moderator level is positive but not significant r = 0.032, t (53) = 0.34, p = 0.739 (Figure 4 B2). rFDG: regional glucose metabolism identified through FDG-PET; CC: clustering coefficient; AD: Alzheimer’s disease; HC: healthy control; HMMR: hierarchical moderated multiple regression; SMA: supplementary motor area; ITG: inferior temporal gyrus.

Demonstrating the fact that AD vs. HC comparison showed a moderating effect of AD on the association between rFDG and changes in CC in these two regions (Supplementary Information), we investigated the association between rFDG and CC in the right ITG and right SMA in each of these groups separately (Figure 4 A2 and B2). In the right ITG there was a negative correlation between rFDG and CC in HC (r = − 0.45, t (53) = − 2.42, p = 0.019), and a positive but non-significant correlation in AD patients (r = 0.1, t (53) = 0.91, p = 0.369) (Figure 4 A2). Similarly in the right SMA, there was an inverse correlation between rFDG and CC in HCs (r = − 0.39, t (53) = − 2.37, p = 0.021), and a trend towards positive correlation in AD patients (r = 0.032, t (53) = 0.34, p = 0.739) (Figure 4 B2).

## DISCUSSION

We examined the association between changes in regional neural activity and topological measures of inter-regional FC in groups of patients with MCI and AD. We demonstrated the following points: *i)* cortical hypometabolism along with disrupted inter-regional FC topology in terms of reduced CC and DC in the regions affected by AD (Supporting Information); *ii)* concordant changes in rFDG and FC topology in form of reduced rFDG associated with reduced CC or DC in the bilateral middle temporal, right angular, right lateral occipital, and right precuneus, as well as the right ITG; *iii)* incongruent changes in form of increased rFDG associated with reduced CC or DC in the bilateral temporal occipital fusiform, bilateral occipital fusiform, and bilateral precentral cortices, as well as the right SMA, in patients with AD and MCI; and *iv)* a significant two-way interaction between rFDG and AD to predict alterations in CC of the right ITG and right SMA. Presence of a significant negative correlation between rFDG and CC in these two areas in healthy controls, and its absence in AD, suggest that AD pathology disrupts the physiologic coupling between rFDG and FC topology.

### The effect of AD on the rFDG-FC topology link

The important question of this study was how could AD affect the physiologic coupling in the association between rFDG and regional topology of functional connectome along the trajectory of disease progression/severity. By simultaneous acquisition of FDG-PET and fMRI data and through a HMMR analysis, we revealed that AD interacts with rFDG to predict regional CC of the right ITG and right SMA. Loss of physiologic correlation between rFDG and CC in patients with AD in our study indicates that AD pathology affects rFDG-CC coupling.

Given the fact that neither ITG, nor SMA are among regions with significant amyloid beta or tau deposition (La Joie, et al., 2012), it is hard to consider the role of AD pathological markers in rFDG-CC decoupling in these regions. As for ITG, significant hypometabolism is seen during early AD, despite minimal amyloid beta deposition (La Joie, et al., 2012; Scheff, et al., 2011). Hypometabolism in the ITG has been reported to be associated with conversion from MCI to AD (Convit, et al., 2000; Morbelli, et al., 2017). This region is involved in verbal fluency and its increased activity predicts better cognitive reserve in AD patients (Halawa, et al., 2019; Weissberger, et al., 2017). Importantly, amyloid beta deposition in signature parietotemporal cortices, is associated with longitudinal tau deposition in the ITG in early AD (Tosun, et al., 2017), highlighting the potential significance of tau pathology over amyloid beta in rFDG/CC decoupling in ITG.

Unlike ITG, neither hypometabolism, nor amyloid depositions are significant in the SMA of patients with AD. Indeed, sparing of glucose metabolism in the primary motor cortex (M1) and SMA has often been reported as a key feature of AD among other types of dementia (Brown, et al., 2014; Marcus, et al., 2014). However, we found higher rFDG activity in the SMA and the precentral gyrus (i.e. where the primary motor or M1 cortex resides) in AD patients. Moreover, reduced task-related FC of the SMA (Cai, et al., 2017; Zheng, et al., 2017), and a paradoxical increase in task-related FC between the SMA and M1 cortex with the sensorimotor, cingulate, and fusiform cortices (Cai, et al., 2017; Vidoni, et al., 2012), i.e. loss of local, and enhanced long-range clustering in functional connectome of the SMA, are in line with reduced CC in the precentral gyrus and SMA of MCI/AD patients in our study.

As indicated earlier, significant amyloid plaque and tau deposition in the M1 and SMA has been reported to occur in stage V and VI of the Braak staging of AD pathology (Braak, et al., 2006; Del Tredici and Braak, 2018; Grothe, et al., 2017; Lyoo, et al., 2016). We suggest that SMA hypermetabolism is a compensatory mechanism to changes that occur in AD pathology as part of system degeneration, rather than direct pathological effect of AD on SMA. This is supported by reported evidence of microstructural amyloid beta and tau aggregation and the resulting neuronal death in the motor cortex of transgenic AD mice before the onset of dementia (Orta-Salazar, et al., 2017).

### The potential role of AD pathology on rFDG and FC topology alterations

Investigating topological changes of the functional connectome in AD has been already reported. Indeed, major functional networks of the brain including the DMN demonstrate profound changes in FC topology in patients with AD and MCI (Dai, et al., 2015; Pereira, et al., 2016; Seo, et al., 2013). The DMN is a major resting state network affected by AD and a site where Aβ deposition, hypometabolism and regional atrophy converge (Buckner, et al., 2008). In particular, reduced efficacy in long-distance functional connections between anterior and posterior parts of the DMN is a prominent feature of AD (Badhwar, et al., 2017; Dillen, et al., 2017; Sanz-Arigita, et al., 2010). The “network degeneration hypothesis” suggests that AD-related pathology initiates in and propagates along specific neuronal populations that follow the trajectory of intrinsic structural or functional brain networks (Palop, et al., 2006; Seeley, et al., 2009; Tahmasian, et al., 2016). Thus, reduced efficacy of long-distance functional connections of the DMN (Liu, et al., 2014) and altered FC topology of hub regions within the posterior DMN and the medial temporal lobe of AD patients (Brier, et al., 2014; Filippi, et al., 2017; Sanz-Arigita, et al., 2010; Seo, et al., 2013; Supekar, et al., 2008), suggest in a “network-based” degeneration pattern for FC topology in AD. Based on this hypothesis it could be speculated that regional AD pathology, namely amyloid, underlies the overlapping patterns of rFDG and FC topology disruption.

Indeed, longitudinal assessments have demonstrated that the severity of amyloid deposition in regions with high degree of amyloid beta pathology (i.e. the posterior DMN), can predict progressive hypometabolism is remote, but functionally connected areas, with minimal amyloid pathology (Klupp, et al., 2014; Klupp, et al., 2015). In another study, rFDG/FC decoupling was found in several areas within target AD regions in the posterior DMN (Marchitelli, et al., 2018), followed by a recent study demonstrating that the rFDG/FC decoupling in the posterior DMN is directly correlated with amyloid beta deposition in this region (Scherr, et al., 2018). In this study, Scherr and colleagues examined the weakening of physiologic coupling between mean rFDG and FC values of the posterior DMN, only to find that rFDG progressively decoupled from FC in the posterior DMN, and the degree of this decoupling associated with mean amyloid beta load in this region (Scherr, et al., 2018). They even identified “rFDG-FC coupling” as the only significant variable that predicted cognitive status of the patients with early and late MCI and AD. We extended their findings here, investigating whether there was a similar physiologic coupling between rFDG and topological metrics of interregional FC outside the DMN in healthy individuals and the influence of MCI/AD on this link.

We adopted CC and DC as more complex measures of FC considering the fact that FC topological measures are shown to be temporally preserved, despite the dynamic nature of resting state FC configurations in health and disease (Liao, et al., 2017). We looked at the whole brain changes in rFDG, CC and DC in MCI and AD patients and identified areas where AD interacted with rFDG to predict changes in CC/DC. We then investigated rFDG-CC coupling in these regions in the control group, and whether it was altered in either of the MCI or AD patients.

Similar to the study by Scherr and colleagues (Scherr, et al., 2018), we found widespread and congruent alterations in rFDG and FC topology in major DMN nodes, including the precuneus, and the middle and inferior temporal gyri, both important sites in DMN, suggesting that Aβ pathology in DMN can be held at least partly accountable for the changes observed in this study.. In addition, we observed rFDG and FC topology alterations in areas outside the DMN, including the lateral occipital cortex, the lingual gyrus and parts of the brain stem (Table 2). We speculated that tau pathology might be responsible for rFDG and topology alterations beyond DMN. This is perhaps supported by the fact that tau deposition is more strongly associated with rFDG decline, than amyloid deposition (Ossenkoppele, et al., 2016). Moreover, connectivity analysis based on tau imaging showed moderate spatial overlap, not only with the dorsal and ventral DMN, but also with visual and language functional networks which comprise the occipital and lingual cortices, respectively (Hoenig, et al., 2018).

Finally, it has been suggested that amyloid beta and tau pathology interact in their regulation of synaptic function. Synaptic tau and amyloid deposition mutually activate precipitating signalling pathways that culminate in progressive synaptic dysfunction and loss. Indeed, it has been demonstrated that in amyloid beta positive normal individuals, functional connectivity has inverse correlation with the degree of tau deposition (Schultz, et al., 2017). A similar model has been described in AD patients, where cascading network failure is mediated by amyloid deposition in the DMN and global tau deposition (Jones, et al., 2017). Based on these findings and our recent model (Pasquini, 2019), we propose that amyloid beta and tau pathology are major driving forces for rFDG and FC topology decline as well as rFDG/FC topology decupling in AD.

Potential drawbacks of this study are its cross-sectional nature of this study and a rather small sample size, which particularly impact drawing a causal relationship between rFDG and topology changes in AD. In addition, we find it imperative to test the pathogenic role of amyloid and tau on metabolism-topology coupling, through simultaneous *in vivo* amyloid and/or tau PET imaging, in the future studies.

## CONCLUSION

This study provides further evidence for network degeneration in AD and extends previous findings of disturbance of functional connectivity in these patients. We showed that there is direct and indirect effect of AD pathology on functional connectivity and its adverse role on physiological coupling between regional metabolism and functional connectivity particularly in the ITG and SMA regions. These findings may further merits investigation using *in vivo* amyloid and/or tau imaging. It would also be useful to evaluate whether amyloid or tau deposition or hypometabolism at the early stages of AD can predict longitudinal alterations in FC topology of signature AD regions, ITG or SMA.

## Acknowledgements

None

## REFERENCES

Aiken, L.S., & West, S. G. (1991) Multiple regression: Testing and interpreting interactions. Thousand Oaks, CA,US. Sage Publications, Inc.

Attwell, D., Laughlin, S.B. (2001) An energy budget for signaling in the grey matter of the brain. Journal of Cerebral Blood Flow & Metabolism, 21:1133–45.

Badhwar, A., Tam, A., Dansereau, C., Orban, P., Hoffstaedter, F., Bellec, P. (2017) Resting-state network dysfunction in Alzheimer’s disease: A systematic review and meta-analysis. Alzheimers Dement (Amst), 8:73–85.

Bischof, G.N., Jessen, F., Fliessbach, K., Dronse, J., Hammes, J., Neumaier, B., Onur, O., Fink, G.R., Kukolja, J., Drzezga, A., van Eimeren, T., Alzheimer’s Disease Neuroimaging, I. (2016) Impact of tau and amyloid burden on glucose metabolism in Alzheimer’s disease. Ann Clin Transl Neurol, 3:934–939.

Braak, H., Alafuzoff, I., Arzberger, T., Kretzschmar, H., Del Tredici, K. (2006) Staging of Alzheimer disease-associated neurofibrillary pathology using paraffin sections and immunocytochemistry. Acta Neuropathol, 112:389–404.

Brier, M.R., Thomas, J.B., Fagan, A.M., Hassenstab, J., Holtzman, D.M., Benzinger, T.L., Morris, J.C., Ances, B.M. (2014) Functional connectivity and graph theory in preclinical Alzheimer’s disease. Neurobiology of aging, 35:757–68.

Brown, R.K., Bohnen, N.I., Wong, K.K., Minoshima, S., Frey, K.A. (2014) Brain PET in suspected dementia: patterns of altered FDG metabolism. Radiographics, 34:684–701.

Buckner, R.L., Andrews-Hanna, J.R., Schacter, D.L. (2008) The brain’s default network: anatomy, function, and relevance to disease. Ann N Y Acad Sci, 1124:1–38.

Bullmore, E., Sporns, O. (2009) Complex brain networks: graph theoretical analysis of structural and functional systems. Nature reviews. Neuroscience, 10:186–98.

Bullmore, E., Sporns, O. (2012) The economy of brain network organization. Nature reviews. Neuroscience, 13:336–49.

Cai, S., Chong, T., Peng, Y., Shen, W., Li, J., von Deneen, K.M., Huang, L. (2017) Altered functional brain networks in amnestic mild cognitive impairment: a resting-state fMRI study. Brain imaging and behavior, 11:619–631.

Chetelat, G., Desgranges, B., Landeau, B., Mezenge, F., Poline, J.B., de la Sayette, V., Viader, F., Eustache, F., Baron, J.C. (2008) Direct voxel-based comparison between grey matter hypometabolism and atrophy in Alzheimer’s disease. Brain, 131:60–71.

Convit, A., de Asis, J., de Leon, M.J., Tarshish, C.Y., De Santi, S., Rusinek, H. (2000) Atrophy of the medial occipitotemporal, inferior, and middle temporal gyri in non-demented elderly predict decline to Alzheimer’s disease. Neurobiology of aging, 21:19–26.

Cranston, I., Reed, L.J., Marsden, P.K., Amiel, S.A. (2001) Changes in regional brain (18)F-fluorodeoxyglucose uptake at hypoglycemia in type 1 diabetic men associated with hypoglycemia unawareness and counter-regulatory failure. Diabetes, 50:2329–36.

Dai, Z., Yan, C., Li, K., Wang, Z., Wang, J., Cao, M., Lin, Q., Shu, N., Xia, M., Bi, Y., He, Y. (2015) Identifying and Mapping Connectivity Patterns of Brain Network Hubs in Alzheimer’s Disease. Cerebral cortex (New York, N.Y. : 1991), 25:3723–42.

Del Tredici, K., Braak, H. (2018) Spreading of Tau Pathology in Sporadic Alzheimer’s Disease Along Cortico-cortical Top-Down Connections. Cerebral Cortex, 28:3372–3384.

Dienel, G.A. (2019) Brain Glucose Metabolism: Integration of Energetics with Function. Physiological reviews, 99:949–1045.

Dillen, K.N.H., Jacobs, H.I.L., Kukolja, J., Richter, N., von Reutern, B., Onur, O.A., Langen, K.J., Fink, G.R. (2017) Functional Disintegration of the Default Mode Network in Prodromal Alzheimer’s Disease. Journal of Alzheimer’s disease : JAD, 59:169–187.

Drzezga, A., Grimmer, T., Henriksen, G., Stangier, I., Perneczky, R., Diehl-Schmid, J., Mathis, C.A., Klunk, W.E., Price, J., DeKosky, S., Wester, H.J., Schwaiger, M., Kurz, A. (2008) Imaging of amyloid plaques and cerebral glucose metabolism in semantic dementia and Alzheimer’s disease. NeuroImage, 39:619–33.

Filippi, M., Basaia, S., Canu, E., Imperiale, F., Meani, A., Caso, F., Magnani, G., Falautano, M., Comi, G., Falini, A., Agosta, F. (2017) Brain network connectivity differs in early-onset neurodegenerative dementia. Neurology, 89:1764–1772.

Friston, K.J., Frith, C.D., Liddle, P.F., Frackowiak, R.S. (1993) Functional connectivity: the principal-component analysis of large (PET) data sets. Journal of Cerebral Blood Flow & Metabolism, 13:5–14.

Gauthier, S., Reisberg, B., Zaudig, M., Petersen, R.C., Ritchie, K., Broich, K., Belleville, S., Brodaty, H., Bennett, D., Chertkow, H., Cummings, J.L., de Leon, M., Feldman, H., Ganguli, M., Hampel, H., Scheltens, P., Tierney, M.C., Whitehouse, P., Winblad, B., International Psychogeriatric Association Expert Conference on mild cognitive, i. (2006) Mild cognitive impairment. Lancet, 367:1262–70.

Gong, G., Rosa-Neto, P., Carbonell, F., Chen, Z.J., He, Y., Evans, A.C. (2009) Age- and gender-related differences in the cortical anatomical network. The Journal of neuroscience : the official journal of the Society for Neuroscience, 29:15684–93.

Greicius, M.D., Srivastava, G., Reiss, A.L., Menon, V. (2004) Default-mode network activity distinguishes Alzheimer’s disease from healthy aging: evidence from functional MRI. Proc Natl Acad Sci U S A, 101:4637–42.

Grothe, M.J., Barthel, H., Sepulcre, J., Dyrba, M., Sabri, O., Teipel, S.J. (2017) In vivo staging of regional amyloid deposition. Neurology, 89:2031–2038.

Halawa, O.A., Gatchel, J.R., Amariglio, R.E., Rentz, D.M., Sperling, R.A., Johnson, K.A., Marshall, G.A. (2019) Inferior and medial temporal tau and cortical amyloid are associated with daily functional impairment in Alzheimer’s disease. Alzheimer’s research & therapy, 11:14.

Hoenig, M.C., Bischof, G.N., Seemiller, J., Hammes, J., Kukolja, J., Onur, O.A., Jessen, F., Fliessbach, K., Neumaier, B., Fink, G.R., van Eimeren, T., Drzezga, A. (2018) Networks of tau distribution in Alzheimer’s disease. Brain : a journal of neurology, 141:568–581.

Jones, D.T., Graff-Radford, J., Lowe, V.J., Wiste, H.J., Gunter, J.L., Senjem, M.L., Botha, H., Kantarci, K., Boeve, B.F., Knopman, D.S., Petersen, R.C., Jack, C.R., Jr. (2017) Tau, amyloid, and cascading network failure across the Alzheimer’s disease spectrum. Cortex; a journal devoted to the study of the nervous system and behavior, 97:143–159.

Kaiser, M. (2011) A tutorial in connectome analysis: topological and spatial features of brain networks. NeuroImage, 57:892–907.

Keuken, M.C., Bazin, P.L., Crown, L., Hootsmans, J., Laufer, A., Muller-Axt, C., Sier, R., van der Putten, E.J., Schafer, A., Turner, R., Forstmann, B.U. (2014) Quantifying inter-individual anatomical variability in the subcortex using 7 T structural MRI. NeuroImage, 94:40–46.

Klupp, E., Forster, S., Grimmer, T., Tahmasian, M., Yakushev, I., Sorg, C., Yousefi, B.H., Drzezga, A. (2014) In Alzheimer’s disease, hypometabolism in low-amyloid brain regions may be a functional consequence of pathologies in connected brain regions. Brain Connect, 4:371–83.

Klupp, E., Grimmer, T., Tahmasian, M., Sorg, C., Yakushev, I., Yousefi, B.H., Drzezga, A., Forster, S. (2015) Prefrontal hypometabolism in Alzheimer disease is related to longitudinal amyloid accumulation in remote brain regions. Journal of nuclear medicine : official publication, Society of Nuclear Medicine, 56:399–404.

La Joie, R., Perrotin, A., Barré, L., Hommet, C., Mézenge, F., Ibazizene, M., Camus, V., Abbas, A., Landeau, B., Guilloteau, D., de La Sayette, V., Eustache, F., Desgranges, B., Chételat, G. (2012) Region-Specific Hierarchy between Atrophy, Hypometabolism, and β-Amyloid (Aβ) Load in Alzheimer&#039;s Disease Dementia. The Journal of Neuroscience, 32:16265.

Latora, V., Marchiori, M. (2001) Efficient behavior of small-world networks. Physical review letters, 87:198701.

Liao, X., Vasilakos, A.V., He, Y. (2017) Small-world human brain networks: Perspectives and challenges. Neuroscience and biobehavioral reviews, 77:286–300.

Liu, Y., Yu, C., Zhang, X., Liu, J., Duan, Y., Alexander-Bloch, A.F., Liu, B., Jiang, T., Bullmore, E. (2014) Impaired long distance functional connectivity and weighted network architecture in Alzheimer’s disease. Cerebral cortex (New York, N.Y. : 1991), 24:1422–35.

Lynall, M.E., Bassett, D.S., Kerwin, R., McKenna, P.J., Kitzbichler, M., Muller, U., Bullmore, E. (2010) Functional connectivity and brain networks in schizophrenia. The Journal of neuroscience : the official journal of the Society for Neuroscience, 30:9477–87.

Lyoo, C.H., Cho, H., Choi, J.Y., Hwang, M.S., Hong, S.K., Kim, Y.J., Ryu, Y.H., Lee, M.S. (2016) Tau Accumulation in Primary Motor Cortex of Variant Alzheimer’s Disease with Spastic Paraparesis. Journal of Alzheimer’s disease : JAD, 51:671–5.

Marchitelli, R., Aiello, M., Cachia, A., Quarantelli, M., Cavaliere, C., Postiglione, A., Tedeschi, G., Montella, P., Milan, G., Salvatore, M., Salvatore, E., Baron, J.C., Pappata, S. (2018) Simultaneous resting-state FDG-PET/fMRI in Alzheimer Disease: Relationship between glucose metabolism and intrinsic activity. NeuroImage, 176:246–258.

Marcus, C., Mena, E., Subramaniam, R.M. (2014) Brain PET in the diagnosis of Alzheimer’s disease. Clinical nuclear medicine, 39:e413–e426.

Masuda, N., Sakaki, M., Ezaki, T., Watanabe, T. (2018) Clustering Coefficients for Correlation Networks. Frontiers in Neuroinformatics, 12.

McKhann, G.M., Knopman, D.S., Chertkow, H., Hyman, B.T., Jack, C.R., Jr., Kawas, C.H., Klunk, W.E., Koroshetz, W.J., Manly, J.J., Mayeux, R., Mohs, R.C., Morris, J.C., Rossor, M.N., Scheltens, P., Carrillo, M.C., Thies, B., Weintraub, S., Phelps, C.H. (2011) The diagnosis of dementia due to Alzheimer’s disease: recommendations from the National Institute on Aging-Alzheimer’s Association workgroups on diagnostic guidelines for Alzheimer’s disease. Alzheimers Dement, 7:263–9.

Meng, C., Brandl, F., Tahmasian, M., Shao, J., Manoliu, A., Scherr, M., Schwerthoffer, D., Bauml, J., Forstl, H., Zimmer, C., Wohlschlager, A.M., Riedl, V., Sorg, C. (2014) Aberrant topology of striatum’s connectivity is associated with the number of episodes in depression. Brain, 137:598–609.

Minoshima, S., Frey, K.A., Koeppe, R.A., Foster, N.L., Kuhl, D.E. (1995) A diagnostic approach in Alzheimer’s disease using three-dimensional stereotactic surface projections of fluorine-18-FDG PET. J Nucl Med, 36:1238–48.

Morbelli, S., Bauckneht, M., Arnaldi, D., Picco, A., Pardini, M., Brugnolo, A., Buschiazzo, A., Pagani, M., Girtler, N., Nieri, A., Chincarini, A., De Carli, F., Sambuceti, G., Nobili, F. (2017) 18F-FDG PET diagnostic and prognostic patterns do not overlap in Alzheimer’s disease (AD) patients at the mild cognitive impairment (MCI) stage. European journal of nuclear medicine and molecular imaging, 44:2073–2083.

Morris, J.C. (1993) The Clinical Dementia Rating (CDR): current version and scoring rules. Neurology, 43:2412–4.

Mosconi, L. (2013) Glucose metabolism in normal aging and Alzheimer’s disease: Methodological and physiological considerations for PET studies. Clinical and translational imaging, 1.

Orta-Salazar, E., Feria-Velasco, A.I., Diaz-Cintra, S. (2017) Primary motor cortex alterations in Alzheimer disease: A study in the 3xTg-AD model. Neurologia (Barcelona, Spain).

Ossenkoppele, R., Schonhaut, D.R., Scholl, M., Lockhart, S.N., Ayakta, N., Baker, S.L., O’Neil, J.P., Janabi, M., Lazaris, A., Cantwell, A., Vogel, J., Santos, M., Miller, Z.A., Bettcher, B.M., Vossel, K.A., Kramer, J.H., Gorno-Tempini, M.L., Miller, B.L., Jagust, W.J., Rabinovici, G.D. (2016) Tau PET patterns mirror clinical and neuroanatomical variability in Alzheimer’s disease. Brain : a journal of neurology, 139:1551–67.

Palop, J.J., Chin, J., Mucke, L. (2006) A network dysfunction perspective on neurodegenerative diseases. Nature, 443:768–73.

Pasquini, L.R., F; Maleki-Balajoo, S; La Joiea, R; Zarei, M; Sorg, C; Drzezga, A; Tahmasian, M.. (2019) Medial temporal lobe disconnection and hyperexcitability across Alzheimer’s disease stages. Journal of Alzheimer’s Disease.

Passow, S., Specht, K., Adamsen, T.C., Biermann, M., Brekke, N., Craven, A.R., Ersland, L., Gruner, R., Kleven-Madsen, N., Kvernenes, O.H., Schwarzlmuller, T., Olesen, R.A., Hugdahl, K. (2015) Default-mode network functional connectivity is closely related to metabolic activity. Hum Brain Mapp, 36:2027–38.

Pereira, J.B., Mijalkov, M., Kakaei, E., Mecocci, P., Vellas, B., Tsolaki, M., Kloszewska, I., Soininen, H., Spenger, C., Lovestone, S., Simmons, A., Wahlund, L.O., Volpe, G., Westman, E. (2016) Disrupted Network Topology in Patients with Stable and Progressive Mild Cognitive Impairment and Alzheimer’s Disease. Cerebral cortex (New York, N.Y. : 1991), 26:3476–3493. project, j. 2018. jamovi (Version 0.9)

Reed, L.J., Marsden, P., Lasserson, D., Sheldon, N., Lewis, P., Stanhope, N., Guinan, E., Kopelman, M.D. (1999) FDG-PET analysis and findings in amnesia resulting from hypoxia. Memory, 7:599–612.

Riedl, V., Bienkowska, K., Strobel, C., Tahmasian, M., Grimmer, T., Forster, S., Friston, K.J., Sorg, C., Drzezga, A. (2014) Local Activity Determines Functional Connectivity in the Resting Human Brain: A Simultaneous FDG-PET/fMRI Study. J Neurosci, 34:6260–6.

Rubinov, M., Sporns, O. (2010) Complex network measures of brain connectivity: uses and interpretations. Neuroimage, 52:1059–69.

Sadeghi, M., Khosrowabadi, R., Bakouie, F., Mahdavi, H., Eslahchi, C., Pouretemad, H. (2017) Screening of autism based on task-free fMRI using graph theoretical approach. Psychiatry Res Neuroimaging, 263:48–56.

Sanz-Arigita, E.J., Schoonheim, M.M., Damoiseaux, J.S., Rombouts, S.A., Maris, E., Barkhof, F., Scheltens, P., Stam, C.J. (2010) Loss of ‘small-world’ networks in Alzheimer’s disease: graph analysis of FMRI resting-state functional connectivity. PLoS One, 5:e13788.

Satterthwaite, T.D., Elliott, M.A., Gerraty, R.T., Ruparel, K., Loughead, J., Calkins, M.E., Eickhoff, S.B., Hakonarson, H., Gur, R.C., Gur, R.E., Wolf, D.H. (2013) An improved framework for confound regression and filtering for control of motion artifact in the preprocessing of resting-state functional connectivity data. Neuroimage, 64:240–56.

Savio, A., Funger, S., Tahmasian, M., Rachakonda, S., Manoliu, A., Sorg, C., Grimmer, T., Calhoun, V., Drzezga, A., Riedl, V., Yakushev, I. (2017) Resting-State Networks as Simultaneously Measured with Functional MRI and PET. Journal of nuclear medicine : official publication, Society of Nuclear Medicine, 58:1314–1317.

Scheff, S.W., Price, D.A., Schmitt, F.A., Scheff, M.A., Mufson, E.J. (2011) Synaptic loss in the inferior temporal gyrus in mild cognitive impairment and Alzheimer’s disease. Journal of Alzheimer’s disease : JAD, 24:547–57.

Scherr, M., Pasquini, L., Benson, G., Nuttall, R., Gruber, M., Neitzel, J., Brandl, F., Sorg, C. (2018) Decoupling of Local Metabolic Activity and Functional Connectivity Links to Amyloid in Alzheimer’s Disease. Journal of Alzheimer’s disease : JAD, 64:405–415.

Scherr, M., Utz, L., Tahmasian, M., Pasquini, L., Grothe, M.J., Rauschecker, J.P., Grimmer, T., Drzezga, A., Sorg, C., Riedl, V. (2019) Effective connectivity in the default mode network is distinctively disrupted in Alzheimer’s disease-A simultaneous resting-state FDG-PET/fMRI study. Hum Brain Mapp.

Schultz, A.P., Chhatwal, J.P., Hedden, T., Mormino, E.C., Hanseeuw, B.J., Sepulcre, J., Huijbers, W., LaPoint, M., Buckley, R.F., Johnson, K.A., Sperling, R.A. (2017) Phases of Hyperconnectivity and Hypoconnectivity in the Default Mode and Salience Networks Track with Amyloid and Tau in Clinically Normal Individuals. The Journal of neuroscience : the official journal of the Society for Neuroscience, 37:4323–4331.

Seeley, W.W., Crawford, R.K., Zhou, J., Miller, B.L., Greicius, M.D. (2009) Neurodegenerative diseases target large-scale human brain networks. Neuron, 62:42–52.

Seo, E.H., Lee, D.Y., Lee, J.M., Park, J.S., Sohn, B.K., Lee, D.S., Choe, Y.M., Woo, J.I. (2013) Whole-brain functional networks in cognitively normal, mild cognitive impairment, and Alzheimer’s disease. PLoS One, 8:e53922.

Sorg, C., Riedl, V., Muhlau, M., Calhoun, V.D., Eichele, T., Laer, L., Drzezga, A., Forstl, H., Kurz, A., Zimmer, C., Wohlschlager, A.M. (2007) Selective changes of resting-state networks in individuals at risk for Alzheimer’s disease. Proc Natl Acad Sci U S A, 104:18760–5.

Sporns, O. (2013) Network attributes for segregation and integration in the human brain. Curr Opin Neurobiol, 23:162–71.

Supekar, K., Menon, V., Rubin, D., Musen, M., Greicius, M.D. (2008) Network analysis of intrinsic functional brain connectivity in Alzheimer’s disease. PLoS computational biology, 4:e1000100.

Tahmasian, M., Pasquini, L., Scherr, M., Meng, C., Forster, S., Mulej Bratec, S., Shi, K., Yakushev, I., Schwaiger, M., Grimmer, T., Diehl-Schmid, J., Riedl, V., Sorg, C., Drzezga, A. (2015) The lower hippocampus global connectivity, the higher its local metabolism in Alzheimer disease. Neurology, 84:1956–63.

Tahmasian, M., Shao, J., Meng, C., Grimmer, T., Diehl-Schmid, J., Yousefi, B.H., Forster, S., Riedl, V., Drzezga, A., Sorg, C. (2016) Based on the Network Degeneration Hypothesis: Separating Individual Patients with Different Neurodegenerative Syndromes in a Preliminary Hybrid PET/MR Study. J Nucl Med, 57:410–5.

Tosun, D., Landau, S., Aisen, P.S., Petersen, R.C., Mintun, M., Jagust, W., Weiner, M.W. (2017) Association between tau deposition and antecedent amyloid-beta accumulation rates in normal and early symptomatic individuals. Brain : a journal of neurology, 140:1499–1512.

van den Heuvel, M.P., Hulshoff Pol, H.E. (2010) Exploring the brain network: a review on resting-state fMRI functional connectivity. Eur Neuropsychopharmacol, 20:519–34.

van Wijk, B.C.M.S., C. J.; Daffertshofer, A. (2010) Comparing Brain Networks of Different Size and Connectivity Density Using Graph Theory. PLoS ONE, 5:e13701.

Vidoni, E.D., Thomas, G.P., Honea, R.A., Loskutova, N., Burns, J.M. (2012) Evidence of altered corticomotor system connectivity in early-stage Alzheimer’s disease. Journal of neurologic physical therapy : JNPT, 36:8–16.

Weissberger, G.H., Melrose, R.J., Fanale, C.M., Veliz, J.V., Sultzer, D.L. (2017) Cortical Metabolic and Cognitive Correlates of Disorientation in Alzheimer’s Disease. Journal of Alzheimer’s disease : JAD, 60:707–719.

Welsh, K.A., Butters, N., Mohs, R.C., Beekly, D., Edland, S., Fillenbaum, G., Heyman, A. (1994) The Consortium to Establish a Registry for Alzheimer’s Disease (CERAD). Part V. A normative study of the neuropsychological battery. Neurology, 44:609–14.

Whisman, M.A., McClelland, G.H. (2005) Designing, testing, and interpreting interactions and moderator effects in family research. J Fam Psychol, 19:111–20.

Yan, C.G., Wang, X.D., Zuo, X.N., Zang, Y.F. (2016) DPABI: Data Processing & Analysis for (Resting-State) Brain Imaging. Neuroinformatics, 14:339–51.

Zheng, W., Liu, X., Song, H., Li, K., Wang, Z. (2017) Altered Functional Connectivity of Cognitive-Related Cerebellar Subregions in Alzheimer’s Disease. Front Aging Neurosci, 9:143.

