## Supplementary file for "Decoupling of inter-regional functional connectivity and regional neural activity in Alzheimer Disease"

1. **Data Acquisition**

**PET images** were started to acquired 30 minutes after injection and recorded for 15 minutes in list-mode acquisition, matrix size = 192$\times$192 pixels, voxel size = 3.7$\times$2.3$\times$2.7 millimeters^3^ (mm^3^), field of view (FOV) = 450 mm and 128 slices (gap = 0.5 mm).

**Structural images:** T1-weighted images were acquired using magnetization-prepared rapid-acquisition gradient echo, repetition time (TR) = 2.300 milliseconds (ms), echo time (TE) = 2.98 ms, FOV = 256 × 256 × 176 mm^3^, voxel size = 1×1×1 mm^3^, flip angle (FA) = 9°, 160 slices (gap = 0.5 mm).

**Functional images:** T2-weighted echo planar imaging (Celle et al.) images were also acquired in interleaved mode with a total scan time of 8 minutes using the following parameters, TR = 2.000 ms, TE = 30 ms, FOV = 192 mm, matrix size = 64×64 pixels, voxel size = 3×3×3 mm^3^, FA = 90°, 35 slices (gap = 0.6 mm).

2. **Preprocessing Imaging data**

Preprocessing of multimodal imaging data started with coregistration of the multimodal imaging dataset to create subject-specific multimodal datasets. Accordingly, raw imaging data were converted to 3D volumes, three volumes of each subject’s functional images were then removed for blood oxygen level dependent (BOLD) signal counterbalance, and the remaining volumes were realigned to the same subject’s mean functional image using a least squares approach with a 6 parameter (rigid body) spatial transformation to estimate head motion during fMRI scan. Excessive head motion (cumulative motion translation or rotation > 3mm or 3°) was applied as an exclusion criterion. Data from one patient with MCI had to be excluded due to excessive movement. In the next step, mean PET image was coregistered to the structural image using rigid-body transformation, followed by both PET and structural images being coregistered to the mean functional image to create subject-specific multimodal datasets.

**Structural and Functional images:** Subject-specific T1-weighted images were segmented to tissue-probability maps of grey matter (GM), white matter (WM) and cerebrospinal fluid (CSF) based on affine regularization procedure. Subject-specific’s EPI and T1-weighted images were then spatially normalized to a standard template provided by the Montreal Neurological Institute (MNI) template using the Statistical Parametric Mapping (SPM12) unified segmentation on T1-weighted routine. MNI tissue-probability maps of gray matter (GM), white matter (WM) and cerebrospinal fluid (CSF) were warped onto single-subject T1-weighted images and stored as masks for later use during the nuisance covariate regression. All EPI images were resampled to an isotropic voxel size of 3×3×3 millimeter (mm). EPI images were smoothed by the use of Gaussian filter (full-width at half-maximum (FWHM) Gaussian kernel of 4 mm). In order to prepare rsfMRI data for functional network analysis, rsfMRI data were also detrended to remove linear trend (Lowe & Russell, 1999). As low frequency (0.01~0.1 Hz) fluctuations of BOLD signal reflects spontaneous neural activity (Lu et al., 2007), bandpass filtering was applied on rsfMRI data to extract signal frequency which are physiological meaningful. Finally, the nuisance variables such as motion parameters with the Frisston-24 model, the WM and the CSF signals, were regressed out to eliminate their effect on BOLD fluctuations (Kelly, Uddin, Biswal, Castellanos, & Milham, 2008). We opted out global signal regression corrections, since this issue is challenging in the rs-fMRI studies (Murphy & Fox, 2017).

**PET images**: Subject-specific PET images were corrected for partial volume effect by the use of PMOD software package (PMOD Technologies Ltd., Adliswil, Switzerland). This software uses the segmented individual T1-weighted image in GM, WM, and CSF for partial volume correction (PVC). PVC-PET images were normalized to the MNI template and resampled to an isotropic voxel size of 3×3×3 mm using SPM12. Finally, PET images were smoothed by the use of Gaussian filter (FWHM Gaussian kernel 12×12×12 mm).

3. **Nodal topological metrics**

To quantify the organization of the whole brain functional connectivity (FC), we used topological metrics calculated using graph theory analyses. These included weighted clustering coefficient (CC) (Saramäki, Kivelä, Onnela, Kaski, & Kertész, 2007) and weighted degree centrality (DC) (Rubinov & Sporns, 2011) metrics, reflecting regional segregation (van den Heuvel & Hulshoff Pol, 2010) and centrality (Sporns, 2013) of FC patterns of brain regions, respectively. Weighted DC represents the sum of the weights of all edges that are directly linked to a node (Rubinov & Sporns, 2011), and is defined as:

$${DC}_{i}^{w}=\sum_{j=1}^{N} w_{ij}$$

where $w_{ij}$ is the weight of connection between node i and j and N is the number of nodes in the graph. The weighted CC for a given node i is equivalent to the fraction of the node neighbors that are also neighbors of each other and is calculated as following:

$${CC}_{i}^{w}=\frac{\sum_{j,h\in N} ({w_{ij}w_{jh}w_{ih})}^{1/3}}{{DC}_{i}^{w}({DC}_{i}^{w}-1)}$$

where ${DC}_{i}^{w}$ is the weighted degree of node i and N is the number of nodes in the graph.

4. **Hierarchical moderated multiple regression (HMMR) analysis**

Starting point in many research projects is to investigate potential associations between variables. However, in some studies the goal is to understand when and how certain associations emerge? This where the question: "under what conditions?" becomes important. The answer to these questions can be addressed by adding a "moderation variable" to the equations. The moderation variable is simply the third variable (M) which has an effect on the link between a predictor variable (X) and outcome variable (Y). The term "moderation effect" is equal to the interaction effect in regression analyses. In other words, a moderator variable changes/interacts with the strength of an association between two other variables. In SI- Figure 1, illustrates two most common forms of moderation effects: conceptual diagram and statistical diagram.

Hierarchical moderated multiple regression (HMMR) analysis is an analytical approach to investigate the effect of moderator variable M on the association between hypothetical variables X and Y which can be formulated as the following:

$$Y=b_{0}+b_{1}X+b_{2}M+b_{3}\left( X*M \right)+e$$

In this model, $b_{0}$ is the intercept of the regression of Y on X that depends on the specific value of M. There is a different line with different slope and intercept for every individual value of M (SI-Figure 2). The moderation (interaction) effect is modeled by the X*M term which is the product of X and M. To form the X*M term, simply multiply together the individual’s scores on X and M. The $b_{3}$ coefficient reflects the interaction between X and M only if the lower order terms $b_{1}X$and $b_{3}M$ are included in the equation**.** To evaluate whether an interaction is significantly present, there are two equivalent ways. 1- we can test whether the coefficient $b_{3}$ differs significantly from zero. 2- we can test whether the increment in the squared multiple correlation (∆R2) given by the interaction is significantly greater than zero. Moreover, we simply add centered continuous covariates as predictors to the regression equation if we want to control for other variables (i.e., covariates of no interest).

5. **Regional neural activity alterations along the trajectory of AD**

Our results displayed a clear pattern of progressive hypometabolism, from HC to MCI and to AD, in the middle temporal, angular, precuneal, and lateral occipital gyri, as well as the ITG (Table 2 and Supplementary Table 1). The middle and inferior temporal, angular and precuneal gyri are all cortical regions of the DMN, showing significant hypometabolism is early AD (La Joie et al., 2012; Morbelli et al., 2017; Scheff, Price, Schmitt, Scheff, & Mufson, 2011). On the other hand, hypometabolism in the lateral occipital gyrus is reported to be minimal in early AD. Lateral occipital hypometabolism is more prominent in lewy body dementia and differentiates these patients from AD (Ishii et al., 1998; Marcus, Mena, & Subramaniam, 2014; Whitwell et al., 2017). Considering the role of this region in object perception (Nagy, Greenlee, & Kovács, 2012), and its functional connection with the DMN during visual object recognition tasks (Karten, Pantazatos, Khalil, Zhang, & Hirsch, 2013), we suggest that lateral occipital hypometabolism might be a heralding sing of the progression of network failure from the DMN onto its functional connections.

Interestingly, our between group rFDG comparisons indicated a progressive hypermetabolism in the temporal occipital fusiform and occipital fusiform cortices in patients with MCI and AD (Table 2 and Supplementary Table 1), which are by definition parts of the visual area 4 of the occipital cortex (Rademacher, Galaburda, Kennedy, Filipek, & Caviness, 1992). The visual cortex shows minimum functional or metabolic decline (Golby et al., 2005; Spehl et al., 2015), and relative sparing from Aβ aggregates in classic forms of AD (La Joie et al., 2012). Indeed, increased rFDG in the fusiform and occipital lobes is associated with cognitive reserve and better performance in visuospatial tasks in AD (Laforce et al., 2014; Matias-Guiu et al., 2017). Therefore, the observed increase in rFDG of occipital regions in AD and MCI patients can be justified in terms of a compensatory hyperactivation in favour of cognitive reserve in these patients. Artificially high metabolism based on the normalisation approach would be an alternative explanation.

6. **Inter-regional FC topology alterations along the trajectory of AD**

AD is associated with a disruption in small-world properties of brains FC topology (Stam, Jones, Nolte, Breakspear, & Scheltens, 2007), in terms of decreased global and local efficacy and increased path lengths (Y. Liu et al., 2014; Z. Liu et al., 2012; Sanz-Arigita et al., 2010). "Small-worldness" is by definition, a property of a graph design where segregation and integration of the connections are both optimized. Small-worldness of FC patterns would therefore interpret as being significantly clustered without reducing the global efficacy (Bullmore & Sporns, 2009). In this regard, a progressive decline in the global efficacy of functional connectomes is observed along the clinical trajectory of AD (E. H. Seo et al., 2013), while alterations in the CC are inconsistent across studies (Brier et al., 2014; Filippi et al., 2017; Sanz-Arigita et al., 2010; E. H. Seo et al., 2013; Supekar, Menon, Rubin, Musen, & Greicius, 2008).

At a regional level, a selective disruption in FC segregation, i.e. reduced CC (Pereira et al., 2016), along with a decrease in nodal centrality, i.e. lower DC, are seen in key AD regions (Caroli et al.; Khazaee, Ebrahimzadeh, Babajani-Feremi, & Alzheimer's Disease Neuroimaging, 2017; Xue & Guo, 2018). Importantly, while reduced regional CC is observed in key AD-affected regions within the medial temporal lobe and posterior DMN (Brier et al., 2014; Filippi et al., 2017; Sanz-Arigita et al., 2010; E. H. Seo et al., 2013; Supekar et al., 2008), whole-brain CC is spared from a similar decrease (E. H. Seo et al., 2013). Instead, global CC is shown to be lowest in MCI patients, and normal or increased in patients with AD (Y. Liu et al., 2014; Z. Liu et al., 2012; Sanz-Arigita et al., 2010; Eun Hyun Seo et al., 2013), suggesting a shift towards a more regular, i.e. clustered, configuration in the functional connectome of AD brain (Liao, Vasilakos, & He, 2017). Combination of the loss of global efficacy and increase in local segregation ultimately result in the so-called "loss of small-worldness" in AD brain. Our findings revealed lower regional CC and DC in AD patients compared to healthy controls, with MCI adopting an intermediate position between AD and HC (supplementary Table 2), in line with the previous findings. Moreover, congruent direction of the alterations in rFDG and CC/DC in most of the significant regions, puts further spin on a common neural basis for these alterations. The important next question is how could AD affect the physiologic coupling of rFDG and regional FC topology.

**SI references:**

Brier, M. R., Thomas, J. B., Fagan, A. M., Hassenstab, J., Holtzman, D. M., Benzinger, T. L., . . . Ances, B. M. (2014). Functional connectivity and graph theory in preclinical Alzheimer's disease. *Neurobiol Aging, 35*(4), 757-768. doi:10.1016/j.neurobiolaging.2013.10.081

Bullmore, E., & Sporns, O. (2009). Complex brain networks: Graph theoretical analysis of structural and functional systems. *Nature Reviews Neuroscience, 10*(3), 186-198. doi:10.1038/nrn2575

Caroli, A., Prestia, A., Chen, K., Ayutyanont, N., Landau, S. M., Madison, C. M., . . . Alzheimer's Disease Neuroimaging, I. (2012). Summary metrics to assess Alzheimer disease-related hypometabolic pattern with 18F-FDG PET: head-to-head comparison. *J Nucl Med, 53*(4), 592-600. doi:10.2967/jnumed.111.094946

Celle, S., Delon-Martin, C., Roche, F., Barthelemy, J. C., Pepin, J. L., & Dojat, M. (2015). Desperately seeking grey matter volume changes in sleep apnea: A methodological review of magnetic resonance brain voxel-based morphometry studies. *Sleep Med Rev*. doi:10.1016/j.smrv.2015.03.001

Filippi, M., Basaia, S., Canu, E., Imperiale, F., Meani, A., Caso, F., . . . Agosta, F. (2017). Brain network connectivity differs in early-onset neurodegenerative dementia. *Neurology, 89*(17), 1764-1772. doi:10.1212/wnl.0000000000004577

Golby, A., Silverberg, G., Race, E., Gabrieli, S., O'Shea, J., Knierim, K., . . . Gabrieli, J. (2005). Memory encoding in Alzheimer's disease: an fMRI study of explicit and implicit memory. *Brain, 128*(Pt 4), 773-787. doi:10.1093/brain/awh400

Ishii, K., Imamura, T., Sasaki, M., Yamaji, S., Sakamoto, S., Kitagaki, H., . . . Mori, E. (1998). Regional cerebral glucose metabolism in dementia with Lewy bodies and Alzheimer&#039;s disease. *Neurology, 51*(1), 125. doi:10.1212/WNL.51.1.125

Karten, A., Pantazatos, S. P., Khalil, D., Zhang, X., & Hirsch, J. (2013). Dynamic coupling between the lateral occipital-cortex, default-mode, and frontoparietal networks during bistable perception. *Brain Connect, 3*(3), 286-293. doi:10.1089/brain.2012.0119

Kelly, A. M., Uddin, L. Q., Biswal, B. B., Castellanos, F. X., & Milham, M. P. (2008). Competition between functional brain networks mediates behavioral variability. *Neuroimage, 39*(1), 527-537. doi:10.1016/j.neuroimage.2007.08.008

Khazaee, A., Ebrahimzadeh, A., Babajani-Feremi, A., & Alzheimer's Disease Neuroimaging, I. (2017). Classification of patients with MCI and AD from healthy controls using directed graph measures of resting-state fMRI. *Behav Brain Res, 322*(Pt B), 339-350. doi:10.1016/j.bbr.2016.06.043

La Joie, R., Perrotin, A., Barré, L., Hommet, C., Mézenge, F., Ibazizene, M., . . . Chételat, G. (2012). Region-Specific Hierarchy between Atrophy, Hypometabolism, and β-Amyloid (Aβ) Load in Alzheimer&#039;s Disease Dementia. *The Journal of Neuroscience, 32*(46), 16265.

Laforce, R., Jr., Tosun, D., Ghosh, P., Lehmann, M., Madison, C. M., Weiner, M. W., . . . Rabinovici, G. D. (2014). Parallel ICA of FDG-PET and PiB-PET in three conditions with underlying Alzheimer's pathology. *Neuroimage Clin, 4*, 508-516. doi:10.1016/j.nicl.2014.03.005

Liao, X., Vasilakos, A. V., & He, Y. (2017). Small-world human brain networks: Perspectives and challenges. *Neurosci Biobehav Rev, 77*, 286-300. doi:10.1016/j.neubiorev.2017.03.018

Liu, Y., Yu, C., Zhang, X., Liu, J., Duan, Y., Alexander-Bloch, A. F., . . . Bullmore, E. (2014). Impaired long distance functional connectivity and weighted network architecture in Alzheimer's disease. *Cereb Cortex, 24*(6), 1422-1435. doi:10.1093/cercor/bhs410

Liu, Z., Zhang, Y., Yan, H., Bai, L., Dai, R., Wei, W., . . . Tian, J. (2012). Altered topological patterns of brain networks in mild cognitive impairment and Alzheimer's disease: a resting-state fMRI study. *Psychiatry Res, 202*(2), 118-125. doi:10.1016/j.pscychresns.2012.03.002

Lowe, M. J., & Russell, D. P. (1999). Treatment of baseline drifts in fMRI time series analysis. *J Comput Assist Tomogr, 23*(3), 463-473.

Lu, H., Zuo, Y., Gu, H., Waltz, J. A., Zhan, W., Scholl, C. A., . . . Stein, E. A. (2007). Synchronized delta oscillations correlate with the resting-state functional MRI signal. *Proc Natl Acad Sci U S A, 104*(46), 18265-18269. doi:10.1073/pnas.0705791104

Marcus, C., Mena, E., & Subramaniam, R. M. (2014). Brain PET in the diagnosis of Alzheimer's disease. *Clin Nucl Med, 39*(10), e413-422; quiz e423-416. doi:10.1097/rlu.0000000000000547

Matias-Guiu, J. A., Cabrera-Martin, M. N., Valles-Salgado, M., Perez-Perez, A., Rognoni, T., Moreno-Ramos, T., . . . Matias-Guiu, J. (2017). Neural Basis of Cognitive Assessment in Alzheimer Disease, Amnestic Mild Cognitive Impairment, and Subjective Memory Complaints. *Am J Geriatr Psychiatry, 25*(7), 730-740. doi:10.1016/j.jagp.2017.02.002

Morbelli, S., Bauckneht, M., Arnaldi, D., Picco, A., Pardini, M., Brugnolo, A., . . . Nobili, F. (2017). 18F-FDG PET diagnostic and prognostic patterns do not overlap in Alzheimer's disease (AD) patients at the mild cognitive impairment (MCI) stage. *Eur J Nucl Med Mol Imaging, 44*(12), 2073-2083. doi:10.1007/s00259-017-3790-5

Murphy, K., & Fox, M. D. (2017). Towards a consensus regarding global signal regression for resting state functional connectivity MRI. *Neuroimage, 154*, 169-173. doi:10.1016/j.neuroimage.2016.11.052

Nagy, K., Greenlee, M. W., & Kovács, G. (2012). The lateral occipital cortex in the face perception network: an effective connectivity study. *Front Psychol, 3*, 141-141. doi:10.3389/fpsyg.2012.00141

Pereira, J. B., Mijalkov, M., Kakaei, E., Mecocci, P., Vellas, B., Tsolaki, M., . . . Westman, E. (2016). Disrupted Network Topology in Patients with Stable and Progressive Mild Cognitive Impairment and Alzheimer's Disease. *Cereb Cortex, 26*(8), 3476-3493. doi:10.1093/cercor/bhw128

Rademacher, J., Galaburda, A. M., Kennedy, D. N., Filipek, P. A., & Caviness, V. S., Jr. (1992). Human cerebral cortex: localization, parcellation, and morphometry with magnetic resonance imaging. *J Cogn Neurosci, 4*(4), 352-374. doi:10.1162/jocn.1992.4.4.352

Rubinov, M., & Sporns, O. (2011). Weight-conserving characterization of complex functional brain networks. *Neuroimage, 56*(4), 2068-2079. doi:10.1016/j.neuroimage.2011.03.069

Sanz-Arigita, E. J., Schoonheim, M. M., Damoiseaux, J. S., Rombouts, S. A., Maris, E., Barkhof, F., . . . Stam, C. J. (2010). Loss of 'small-world' networks in Alzheimer's disease: graph analysis of FMRI resting-state functional connectivity. *PLOS ONE, 5*(11), e13788. doi:10.1371/journal.pone.0013788

Saramäki, J., Kivelä, M., Onnela, J. P., Kaski, K., & Kertész, J. (2007). Generalizations of the clustering coefficient to weighted complex networks.

. *Physical review. E, Statistical, nonlinear, and soft matter physics, E 75*, 027105.

Scheff, S. W., Price, D. A., Schmitt, F. A., Scheff, M. A., & Mufson, E. J. (2011). Synaptic loss in the inferior temporal gyrus in mild cognitive impairment and Alzheimer's disease. *J Alzheimers Dis, 24*(3), 547-557. doi:10.3233/jad-2011-101782

Seo, E. H., Lee, D. Y., Lee, J.-M., Park, J.-S., Sohn, B. K., Lee, D. S., . . . Woo, J. I. (2013). Whole-brain Functional Networks in Cognitively Normal, Mild Cognitive Impairment, and Alzheimer’s Disease. *PLOS ONE, 8*(1), e53922. doi:10.1371/journal.pone.0053922

Seo, E. H., Lee, D. Y., Lee, J. M., Park, J. S., Sohn, B. K., Lee, D. S., . . . Woo, J. I. (2013). Whole-brain functional networks in cognitively normal, mild cognitive impairment, and Alzheimer's disease. *PLoS One, 8*(1), e53922. doi:10.1371/journal.pone.0053922

Spehl, T. S., Hellwig, S., Amtage, F., Weiller, C., Bormann, T., Weber, W. A., . . . Frings, L. (2015). Syndrome-specific patterns of regional cerebral glucose metabolism in posterior cortical atrophy in comparison to dementia with Lewy bodies and Alzheimer's disease--a [F-18]-FDG pet study. *J Neuroimaging, 25*(2), 281-288. doi:10.1111/jon.12104

Sporns, O. (2013). Network attributes for segregation and integration in the human brain. *Curr Opin Neurobiol, 23*(2), 162-171. doi:10.1016/j.conb.2012.11.015

Stam, C. J., Jones, B. F., Nolte, G., Breakspear, M., & Scheltens, P. (2007). Small-world networks and functional connectivity in Alzheimer's disease. *Cereb Cortex, 17*(1), 92-99. doi:10.1093/cercor/bhj127

Supekar, K., Menon, V., Rubin, D., Musen, M., & Greicius, M. D. (2008). Network analysis of intrinsic functional brain connectivity in Alzheimer's disease. *PLoS Comput Biol, 4*(6), e1000100. doi:10.1371/journal.pcbi.1000100

van den Heuvel, M. P., & Hulshoff Pol, H. E. (2010). Exploring the brain network: a review on resting-state fMRI functional connectivity. *Eur Neuropsychopharmacol, 20*(8), 519-534. doi:10.1016/j.euroneuro.2010.03.008

Whitwell, J. L., Graff-Radford, J., Singh, T. D., Drubach, D. A., Senjem, M. L., Spychalla, A. J., . . . Josephs, K. A. (2017). (18)F-FDG PET in Posterior Cortical Atrophy and Dementia with Lewy Bodies. *J Nucl Med, 58*(4), 632-638. doi:10.2967/jnumed.116.179903

Xue, S. W., & Guo, Y. (2018). Increased resting-state brain entropy in Alzheimer's disease. *Neuroreport, 29*(4), 286-290. doi:10.1097/wnr.0000000000000942

**SI-TABLE 1- Analysis of variance on regional glucose metabolism for each region in Harvard-Oxford Atlas. Post-hoc test: permutation test (p<0.05, 100,000 permutations).**

| **Brain Regions** | **HC** | **MCI** | **AD** | **Post-hoc** | |
| --- | --- | --- | --- | --- | --- |
|  | 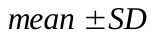 | 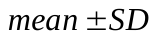 | 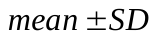 | **Group comparison** | **p-value** |
| **Precentral Gyrus (L)** | 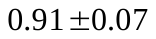 | 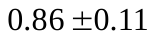 | 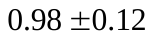 | HC>MCI  HC<AD  MCI<AD | 0.04  0.002  0.0002 |
| **Precentral Gyrus (R)** | 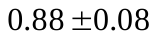 | 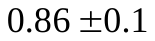 | 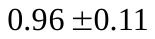 | HC<AD  MCI<AD | 0.001  <0.0001 |
| **Middle Temporal Gyrus, posterior (L)** | 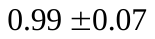 | 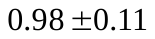 | 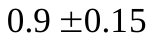 | HC>AD  MCI>AD | 0.002  0.02 |
| **Middle Temporal Gyrus, temporooccipital (L)** | 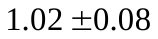 | 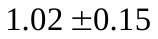 | 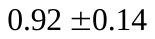 | HC>AD  MCI>AD | 0.001  0.01 |
| **Inferior Temporal Gyrus, posterior (L)** | 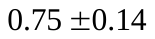 | 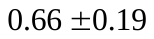 | 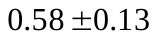 | HC>MCI  HC>AD  MCI>AD | 0.04  0  0.03 |
| **Inferior Temporal Gyrus, posterior (R)** | 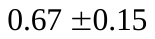 | 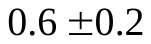 | 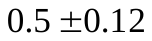 | HC>AD  MCI>AD | <0.0001  0.049 |
| **Inferior Temporal Gyrus, temporooccipital (L)** | 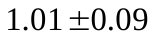 | 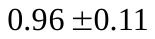 | 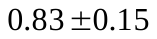 | HC>MCI  HC>AD  MCI>AD | 0.048  0  0.0002 |
| **Inferior Temporal Gyrus, temporooccipital(R)** | 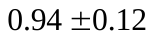 | 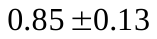 | 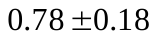 | HC>MCI  HC>AD | 0.006  <0.0001 |
| **Angular Gyrus (L)** | 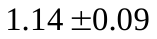 | 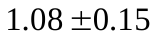 | 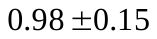 | HC>MCI  HC>AD  MCI>AD | 0.04  <0.0001  0.01 |
| **Angular Gyrus (R)** | 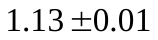 | 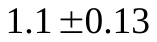 | 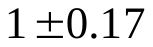 | HC>AD  MCI>AD | <0.0001  0.01 |
| **Lateral Occipital Cortex, superior (L)** | 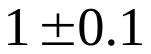 | 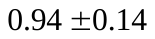 | 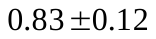 | HC>AD  MCI>AD | 0  0.003 |
| **Lateral Occipital Cortex, superior (R)** | 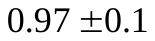 | 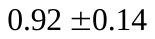 | 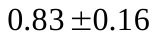 | HC>AD  MCI>AD | <0.0001  0.02 |
| **Supplementary Motor Area (R)** | 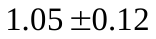 | 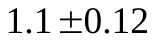 | 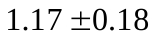 | HC<AD | 0.001 |
| **Precuneous Cortex (L)** | 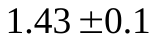 | 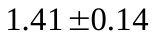 | 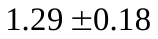 | HC>AD  MCI>AD | 0.006  0.01 |
| **Precuneous Cortex (R)** | 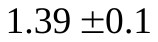 | 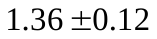 | 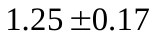 | HC>AD  MCI>AD | 0.006  0.01 |
| **Parahippocampal Gyrus, posterior (L)** | 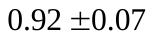 | 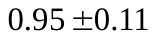 | 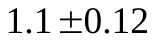 | HC<AD  MCI<AD | 0.0004  0.04 |
| **Parahippocampal Gyrus, posterior (R)** | 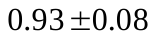 | 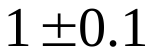 | 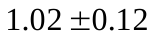 | HC<MCI  HC<AD | 0.006  0.001 |
| **Lingual Gyrus (L)** | 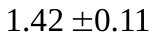 | 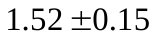 | 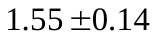 | HC<MCI  HC<AD | 0.007  0.0001 |
| **Lingual Gyrus (R)** | 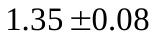 | 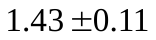 | 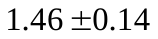 | HC<MCI  HC<AD | 0.004  0.0004 |
| **Temporal Occipital Fusiform Cortex (L)** | 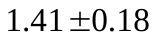 | 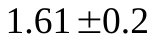 | 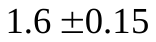 | HC<MCI  HC<AD | 0.0002  0 |
| **Temporal Occipital Fusiform Cortex (R)** | 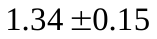 | 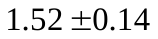 | 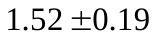 | HC<MCI  HC<AD | <0.0001  <0.0001 |
| **Occipital Fusiform Gyrus (L)** | 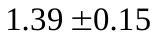 | 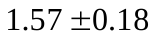 | 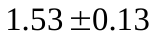 | HC<MCI  HC<AD | <0.0001  0.0001 |
| **Occipital Fusiform Gyrus (R)** | 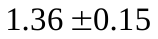 | 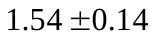 | 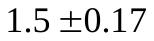 | HC<MCI  HC<AD | <0.0001  0.006 |
| **Brain-Stem(L)** | 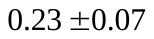 | 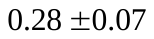 | 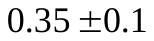 | HC<MCI  HC<AD  MCI<AD | 0.01  0  0.005 |
| **Brain-Stem(R)** | 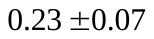 | 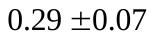 | 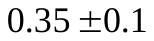 | HC<MCI  HC<AD  MCI<AD | 0.007  0  0.009 |
| **Putamen (L)** | 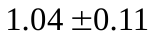 | 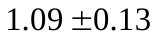 | 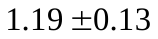 | HC<AD  MCI<AD | 0  0.003 |
| **Hippocampus (R)** | 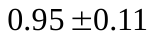 | 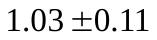 | 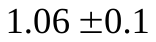 | HC<MCI  HC<AD | 0.009  <0.0001 |
| **Amygdala (R)** | 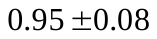 | 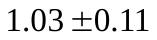 | 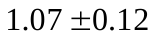 | HC<MCI  HC<AD | 0.008  <0.0001 |
| **Accumbens (R)** | 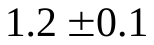 | 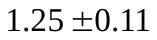 | 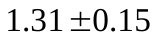 | HC<AD | 0.001 |

AD: Alzheimer disease; HC: healthy controls; MCI: mild cognitive impairment; SD: standard deviation; L: left; R: right.

**SI-TABLE 2- Analysis of variance on clustering coefficient as a nodal metric for each region in Harvard-Oxford Atlas. Post-hoc test: permutation test (p<0.05, 100,000 permutations).**

| **Brain Regions** | **HC** | **MCI** | **AD** | **Post-hoc** | |
| --- | --- | --- | --- | --- | --- |
|  | 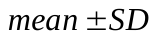 | 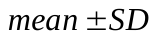 | 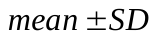 | **Group comparison** | **p-value** |
| **Frontal Pole (L)** | 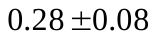 | 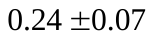 | 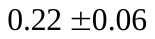 | HC>AD | 0.001 |
| **Frontal Pole (R)** | 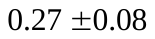 | 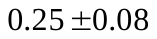 |  | HC>AD  MCI>AD | 0.0005  0.008 |
| **Insular Cortex (L)** |  |  |  | HC>MCI  HC>AD | 0.02  0.0001 |
| **Insular Cortex (R)** |  |  |  | HC>MCI  HC>AD  MCI>AD | 0.04  0  0.004 |
| **Superior Frontal Gyrus (L)** |  |  |  | HC>AD  MCI>AD | 0.0006  0.03 |
| **Middle Frontal Gyrus (L)** |  |  |  | HC>AD  MCI>AD | <0.0001  0.007 |
| **Middle Frontal Gyrus (R)** |  |  |  | HC>AD  MCI>AD | <0.0001  0.004 |
| **Inferior Frontal Gyrus, pars triangularis (L)** |  |  |  | HC>AD  MCI>AD | 0  0.003 |
| **Inferior Frontal Gyrus, pars triangularis (R)** |  |  |  | HC>AD  MCI>AD | <0.0001  0.007 |
| **Inferior Frontal Gyrus, pars opercularis (L)** |  |  |  | HC>AD  MCI>AD | 0.0004  0.008 |
| **Inferior Frontal Gyrus, pars opercularis (R)** |  |  |  | HC>MCI  HC>AD  MCI>AD | 0.03  0  0.02 |
| **Precentral Gyrus (L)** |  |  |  | HC>MCI  HC>AD | 0.004  <0.0001 |
| **Precentral Gyrus (R)** |  |  |  | HC>MCI  HC>AD  MCI>AD | 0.008  <0.0001  0.047 |
| **Temporal Pole (L)** |  |  |  | HC>MCI  HC>AD | 0.018  0.0004 |
| **Temporal Pole (R)** |  |  |  | HC>AD  MCI>AD | <0.0001  0.009 |
| **Superior Temporal Gyrus, anterior (L)** |  |  |  | HC>MCI  HC>AD | 0.005  0.0001 |
| **Superior Temporal Gyrus, anterior (R)** |  |  |  | HC>MCI  HC>AD | 0.024  0.001 |
| **Superior Temporal Gyrus, posterior (L)** |  |  |  | HC>MCI  HC>AD  MCI>AD | 0.04  0  0.003 |
| **Superior Temporal Gyrus, posterior (R)** |  |  |  | HC>MCI  HC>AD  MCI>AD | 0.007  0  0.14 |
| **Middle Temporal Gyrus, anterior (R)** |  |  |  | HC>MCI  HC>AD | 0.009  <0.0001 |
| **Middle Temporal Gyrus, posterior (L)** |  |  |  | HC>AD  MCI>AD | 0.002  0.005 |
| **Middle Temporal Gyrus, posterior (R)** |  |  |  | HC>AD  MCI>AD | 0.0001  0.02 |
| **Middle Temporal Gyrus, temporooccipital (L)** |  |  |  | HC>MCI  HC>AD | 0.03  0.0003 |
| **Middle Temporal Gyrus, temporooccipital (R)** |  |  |  | HC>MCI  HC>AD | 0.02  0.0003 |
| **Inferior Temporal Gyrus, temporooccipital (L)** |  |  |  | HC>MCI  HC>AD | 0.01  0.0001 |
| **Inferior Temporal Gyrus, temporooccipital (R)** |  |  |  | HC>AD  MCI>AD | 0.0002  0.02 |
| **Postcentral Gyrus (L)** |  |  |  | HC>MCI  HC>AD  MCI>AD | 0.009  0  0.01 |
| **Postcentral Gyrus (R)** |  |  |  | HC>AD  MCI>AD | 0  0.003 |
| **Superior Parietal Lobule (L)** |  |  |  | HC>MCI  HC>AD | 0.003  0 |
| **Superior Parietal Lobule (R)** |  |  |  | HC>MCI  HC>AD | 0.003  0 |
| **Supramarginal Gyrus, anterior (L)** |  |  |  | HC>MCI  HC>AD  MCI>AD | 0.01  <0.0001  0.03 |
| **Supramarginal Gyrus, anterior (R)** |  |  |  | HC>MCI  HC>AD  MCI>AD | 0.03  <0.0001  0.02 |
| **Supramarginal Gyrus, posterior (R)** |  |  |  | HC>MCI  HC>AD  MCI>AD | 0.02  <0.0001  0.03 |
| **Angular Gyrus (R)** |  |  |  | HC>AD  MCI>AD | 0.002  0.02 |
| **Lateral Occipital Cortex, superior (R)** |  |  |  | HC>AD  MCI>AD | <0.0001  0.02 |
| **Lateral Occipital Cortex, inferior (L)** |  |  |  | HC>MCI  HC>AD | 0.04  0.0009 |
| **Lateral Occipital Cortex, inferior (R)** |  |  |  | HC>AD  MCI>AD | <0.0001  0.004 |
| **Frontal Medial Cortex (L)** |  |  |  | HC>AD  MCI>AD | 0.0002  0.04 |
| **Frontal Medial Cortex (R)** |  |  |  | HC>AD  MCI>AD | 0.0001  0.02 |
| **Supplementary Motor Area (L)** |  |  |  | HC>MCI  HC>AD  MCI>AD | 0.02  <0.0001  0.03 |
| **Supplementary Motor Area (R)** |  |  |  | HC>MCI  HC>AD  MCI>AD | 0.001  0  0.003 |
| **Cingulate Gyrus, anterior (L)** |  |  |  | HC>AD  MCI>AD | 0.0004  0.04 |
| **Cingulate Gyrus, posterior (L)** |  |  |  | HC>AD  MCI>AD | 0.0004  0.03 |
| **Cingulate Gyrus, posterior (R)** |  |  |  | HC>MCI  HC>AD  MCI>AD | 0.02  <0.0001  0.049 |
| **Precuneous Cortex (R)** |  |  |  | HC>MCI  HC>AD | 0.02  0.0005 |
| **Cuneal Cortex (L)** |  |  |  | HC>AD  MCI >AD | 0.0006  0.04 |
| **Frontal Orbital Cortex (R)** |  |  |  | HC>AD  MCI >AD | <0.0001  0.007 |
| **Temporal Occipital Fusiform Cortex (L)** |  |  |  | HC>MCI  HC>AD | 0.03  0.002 |
| **Temporal Occipital Fusiform Cortex (R)** |  |  |  | HC>MCI  HC>AD | 0.003  0 |
| **Occipital Fusiform Gyrus (L)** |  |  |  | HC>MCI  HC>AD | 0.009  0.0001 |
| **Occipital Fusiform Gyrus (R)** |  |  |  | HC>MCI  HC>AD  MCI>AD | 0.02  <0.0001  0.01 |
| **Frontal Operculum Cortex (L)** |  |  |  | HC>MCI  HC>AD | 0.03  0.001 |
| **Frontal Operculum Cortex (R)** |  |  |  | HC>AD  MCI >AD | 0  0.003 |
| **Central Opercular Cortex (L)** |  |  |  | HC>MCI  HC>AD  MCI>AD | 0.003  0  0.01 |
| **Central Opercular Cortex (R)** |  |  |  | HC>MCI  HC>AD  MCI>AD | 0.002  0  0.048 |
| **Parietal Operculum Cortex (L)** |  |  |  | HC>MCI  HC>AD | 0.002  0 |
| **Parietal Operculum Cortex (R)** |  |  |  | HC>MCI  HC>AD  MCI>AD | 0.02  0  0.03 |
| **PlanumPolare (L)** |  |  |  | HC>MCI  HC>AD  MCI>AD | 0.02  <0.0001  0.01 |
| **PlanumPolare (R)** |  |  |  | HC>MCI  HC>AD  MCI>AD | 0.03  <0.0001  0.01 |
| **Heschls Gyrus (includes H1 and H2) (L)** |  |  |  | HC>MCI  HC>AD  MCI>AD | 0.03  0  0.02 |
| **Heschls Gyrus (includes H1 and H2) (R)** |  |  |  | HC>MCI  HC>AD  MCI>AD | 0.001  0  0.03 |
| **PlanumTemporale (L)** |  |  |  | HC>MCI  HC>AD  MCI>AD | 0.008  0  0.04 |
| **PlanumTemporale (R)** |  |  |  | HC>MCI  HC>AD  MCI>AD | 0.006  0  0.03 |
| **Occipital Pole (L)** |  |  |  | HC>AD  MCI>AD | 0.0005  0.01 |
| **Occipital Pole (R)** |  |  |  | HC>AD | 0.0008 |
| **Hippocampus (L)** |  |  |  | HC>MCI  HC>AD  MCI>AD | 0.044  0.0003  0.03 |

AD: Alzheimer disease; HC: healthy controls; MCI: mild cognitive impairment; SD: standard deviation; L: left; R: right.

**SI-TABLE 3- Analysis of variance on degree centrality as a nodal metric for each region in Harvard-Oxford Atlas. Post-hoc test: permutation test (p<0.05, 100,000 permutations).**

| **Brain Regions** | **HC** | **MCI** | **AD** | **Post-hoc** | |
| --- | --- | --- | --- | --- | --- |
|  |  |  |  | **Group comparison** | **p-value** |
| **Insular Cortex (L)** |  |  |  | HC>MCI  HC>AD | 0.003  0.0003 |
| **Insular Cortex (R)** |  |  |  | HC>MCI  HC>AD | 0.0006  <0.0001 |
| **Inferior Frontal Gyrus, pars opercularis (R)** |  |  |  | HC>AD  MCI>AD | 0.0005  0.02 |
| **Precentral Gyrus (L)** |  |  |  | HC>AD  MCI>AD | 0.0002  0.02 |
| **Precentral Gyrus (R)** |  |  |  | HC>MCI  HC>AD  MCI>AD | 0.03  0.0003  0.046 |
| **Temporal Pole (R)** |  |  |  | HC>MCI  HC>AD | 0.01  0.0002 |
| **Superior Temporal Gyrus, posterior (R)** |  |  |  | HC>MCI  HC>AD | 0.01  <0.0001 |
| **Postcentral Gyrus (L)** |  |  |  | HC>MCI  HC>AD | 0.03  0.002 |
| **Postcentral Gyrus (R)** |  |  |  | HC>MCI  HC>AD | 0.0006  0.0004 |
| **Superior Parietal Lobule (L)** |  |  |  | HC>MCI  HC>AD | 0.01  0.0009 |
| **Superior Parietal Lobule (R)** |  |  |  | HC>MCI  HC>AD | 0.006  <0.0001 |
| **Lateral Occipital Cortex, superior (R)** |  |  |  | HC>MCI  HC>AD | 0.007  0.0006 |
| **Lateral Occipital Cortex, inferior (L)** |  |  |  | HC>MCI  HC>AD | 0.004  0.0004 |
| **Lateral Occipital Cortex, inferior (R)** |  |  |  | HC>MCI  HC>AD | 0.001  0.002 |
| **Supplementary Motor Area (L)** |  |  |  | HC>MCI  HC>AD | 0.02  <0.0001 |
| **Cuneal Cortex (L)** |  |  |  | HC>MCI  HC>AD | 0.005  0.002 |
| **Cuneal Cortex (R)** |  |  |  | HC>MCI  HC>AD | 0.0006  0.0003 |
| **Temporal Fusiform Cortex, anterior (L)** |  |  |  | HC<MCI  HC<AD | 0.009  0.001 |
| **Temporal Fusiform Cortex, anterior (R)** |  |  |  | HC<MCI  HC<AD | 0.005  0.0008 |
| **Central Opercular Cortex (L)** |  |  |  | HC>MCI  HC>AD | 0.02  0.0003 |
| **Central Opercular Cortex (R)** |  |  |  | HC>MCI  HC>AD | 0.005  <0.0001 |
| **Parietal Operculum Cortex (L)** |  |  |  | HC>MCI  HC>AD  MCI>AD | 0.03  <0.0001  0.04 |
| **Parietal Operculum Cortex (R)** |  |  |  | HC>MCI  HC>AD | 0.0003  <0.0001 |
| **PlanumPolare (R)** |  |  |  | HC>MCI  HC>AD | 0.001  <0.0001 |
| **Heschls Gyrus (includes H1 and H2) (L)** |  |  |  | HC>MCI  HC>AD | 0.009  <0.0001 |
| **Heschls Gyrus (includes H1 and H2) (R)** |  |  |  | HC>MCI  HC>AD | 0.03  0.0004 |
| **PlanumTemporale (L)** |  |  |  | HC>MCI  HC>AD  MCI>AD | 0.042  <0.0001  0.04 |
| **PlanumTemporale (R)** |  |  |  | HC>MCI  HC>AD  MCI>AD | 0.02  0  0.02 |
| **Supracalcarine Cortex (L)** |  |  |  | HC>MCI  HC>AD | 0.007  0.004 |
| **Supracalcarine Cortex (R)** |  |  |  | HC>MCI  HC>AD | 0.02  0.002 |
| **Brain-Stem(R)** |  |  |  | HC<AD  MCI<AD | 0.008  0.0005 |
| **Pallidum(L)** |  |  |  | HC<MCI  HC<AD | 0.006  0.0003 |

AD: Alzheimer disease; HC: healthy controls; MCI: mild cognitive impairment; SD: standard deviation; L: left; R: right.

**SI-Figures**

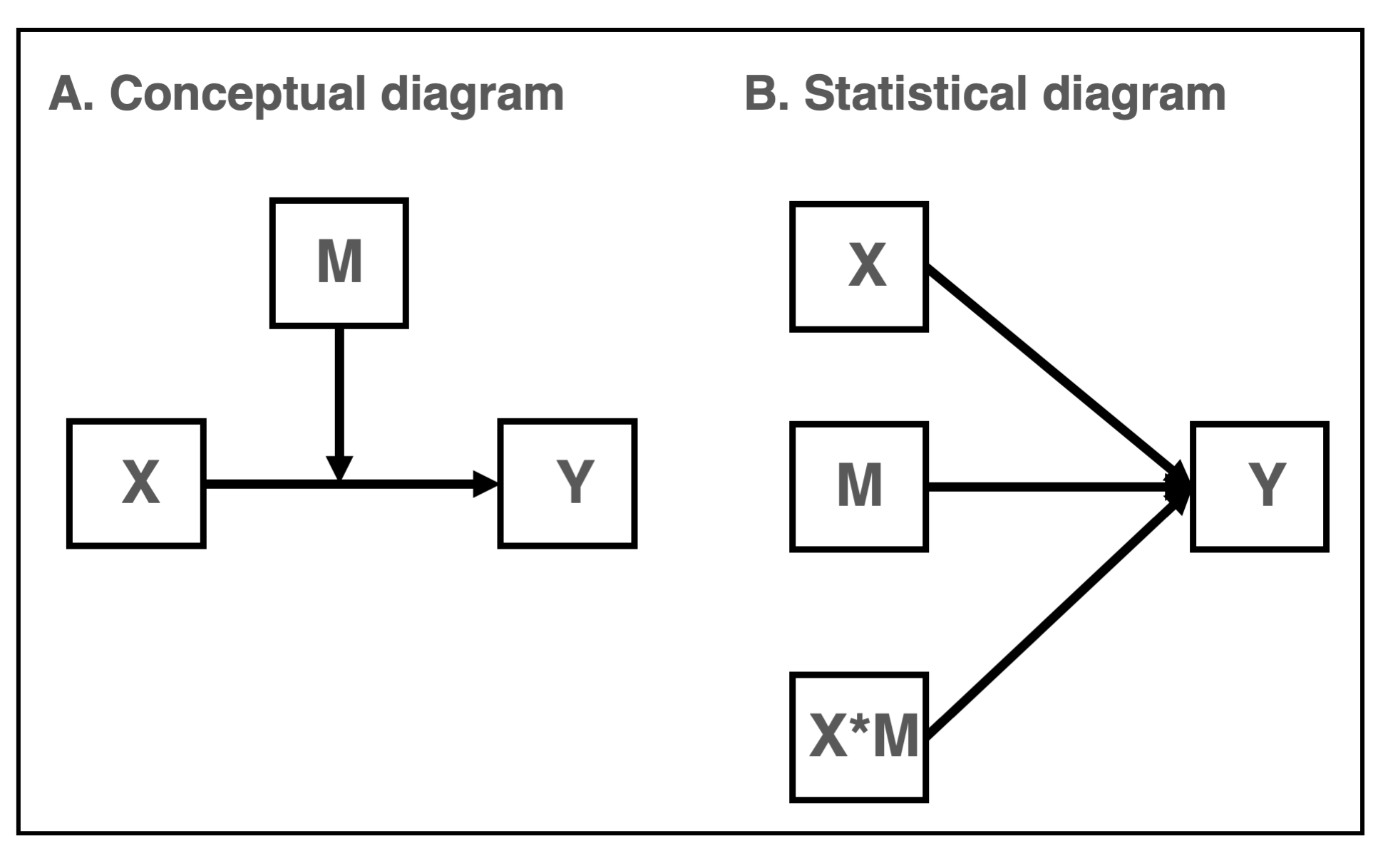

**SI-Figure 1**: Schematic diagrams of moderation are illustrated as A) conceptual diagram and B) statistical diagram.

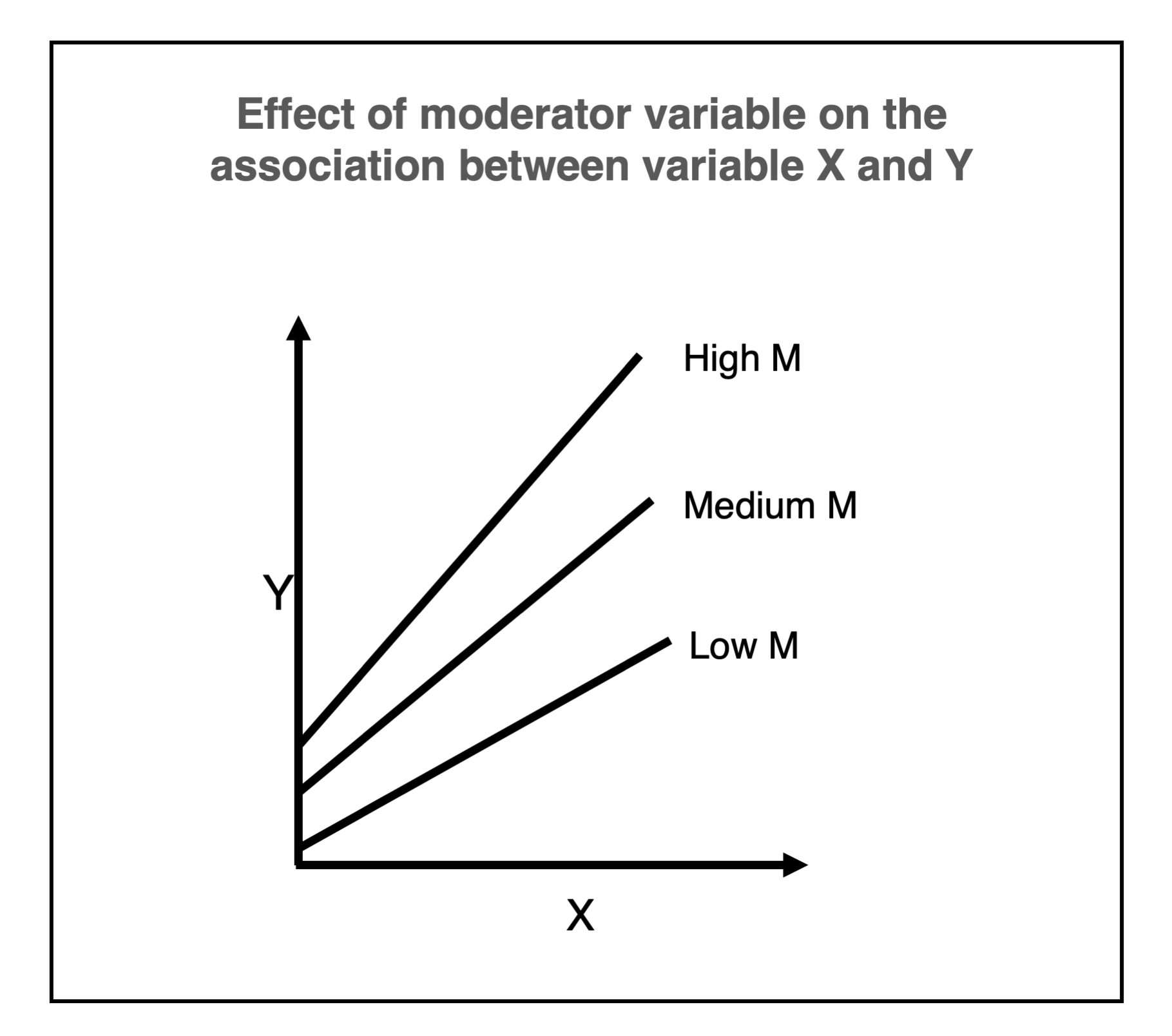

**SI-Figure 2**: The effect of moderator variable on the association between variable X and Y. The slop and intercept of the regression of Y on X that depends on the specific value of M (High, Medium, or Low).
